# Synapse-selective control of cortical maturation and plasticity engages an interneuron-autonomous synaptic switch

**DOI:** 10.1101/382614

**Authors:** Adema Ribic, Michael C. Crair, Thomas Biederer

## Abstract

**Highlights:**

- The synaptogenic molecule SynCAM 1 is selectively regulated by visual experience
- SynCAM 1 controls thalamic input onto cortical Parvalbumin (PV^+^) interneurons
- PV^+^-specific knockdown of SynCAM 1 arrests maturation of cortical inhibition
- Thalamic excitation onto PV^+^ interneurons is essential for critical period closure

**eTOC Blurb:** Ribic et al. show that network plasticity in both young and adult cortex is restricted by the synapse organizing molecule SynCAM 1. On a cellular level, it functions in Parvalbumin-positive interneurons to recruit thalamocortical terminals. This controls the maturation of inhibitory drive and restricts plasticity in the cortex. These results reveal the synaptic locus of cortical plasticity and identify the first cell-autonomous synaptic factor for closure of cortical critical periods.

**Summary:** Brain plasticity peaks early in life and tapers in adulthood. This is exemplified in the primary visual cortex, where brief loss of vision to one eye abrogates cortical responses to inputs from that eye during the critical period, but not in adulthood. The synaptic locus of critical period plasticity and the cell-autonomous synaptic factors timing these periods remain unclear. We here demonstrate that the immunoglobulin protein Synaptic Cell Adhesion Molecule 1 (SynCAM 1/Cadm1) is regulated by visual experience and limits visual cortex plasticity. SynCAM 1 selectively controls the number of excitatory thalamocortical (TC) inputs onto Parvalbumin (PV^+^) interneurons and loss of SynCAM 1 in turn impairs the maturation of TC-driven feed-forward inhibition. SynCAM 1 acts in cortical PV^+^ interneurons to perform these functions and its PV^+^-specific knockdown prevents the age-related plasticity decline. These results identify a synapse type-specific, cell-autonomous mechanism that governs circuit maturation and closes the visual critical period.

## Introduction

Sensory input during developmental critical periods shapes neural circuits and drives their functional maturation (Hensch, 2005; Levelt and Hubener, 2012; Wang et al., 2010). The quality of sensory experience is a critical factor in the development of circuit function. Imbalanced visual input during the critical period for vision in early human childhood (up to 8 years of age; Hooks and Chen, 2007) leads to a permanent deficit in vision called amblyopia (Espinosa and Stryker, 2012; Hubener and Bonhoeffer, 2014). Experimentally, occlusion of one eye during the critical period for vision in cats (postnatal weeks 4-8; Hubel and Wiesel, 1970) and mice (postnatal days 25-32; Frenkel and Bear, 2004; Gordon and Stryker, 1996) leads to a loss of cortical responses to the closed eye and an increase in responses to the open eye, a phenomenon known as ocular dominance plasticity (ODP). The elevated potential for ODP during the critical period facilitates the extensive experience-dependent structural and functional refinement of synapses that takes place during early postnatal development of the cortex (Espinosa and Stryker, 2012; Hooks and Chen, 2007; Wang et al., 2010). Plasticity tapers off as the brain matures, such that short visual deprivation in adult animals has no effect on cortical responses (Lehmann and Lowel, 2008).

ODP is traditionally viewed as a cortical phenomenon (Espinosa and Stryker, 2012; Kuhlman et al., 2013), but the precise synaptic loci of critical period plasticity remain to be determined due to complex feed-forward and feedback connectivity between sensory thalamus and cortex (Jaepel et al., 2017; Sommeijer et al., 2017; Thompson et al., 2016). There is considerable evidence that the maturation of cortical inhibition controls the opening of the critical period for vision (Fagiolini and Hensch, 2000; Kuhlman et al., 2013; Southwell et al., 2010). Robust activation of Parvalbumin (PV^+^) interneurons by thalamocortical (TC) axons at the onset of sensory activity drives the development of cortical inhibition (Chittajallu and Isaac, 2010; Cruikshank et al., 2007; Shen and Colonnese, 2016). TC axons continue to activate PV^+^ interneurons more strongly than pyramidal neurons, maintaining the high inhibitory drive in the mature brain (Chittajallu and Isaac, 2010; Cruikshank et al., 2007; Ji et al., 2016; Kloc and Maffei, 2014). Reducing this inhibitory drive can extend plasticity beyond the critical period and even reactivate it in adult animals (Harauzov et al., 2010; Kuhlman et al., 2013), and transplantation of inhibitory neurons to adult cortex can create a new critical period (Southwell et al., 2010). Removal of so-called molecular brakes delays critical period closure (Takesian and Hensch, 2013). Known molecular brakes include extracellular matrix (ECM) components (Pizzorusso et al., 2002), cholinergic signaling (Morishita et al., 2010), epigenetic factors (Nott et al., 2015) and myelin-associated receptors NogoR and PirB (McGee et al., 2005; Syken et al., 2006). Apart from PirB, a pyramidal neuron-expressed receptor for myelin-associated axonal growth inhibitors (Atwal et al., 2008; Vidal et al., 2016), recent research suggests that molecular brake mechanisms converge on cortical inhibitory, fast spiking Parvalbumin (PV^+^) cells (Demars and Morishita, 2014; Lensjo et al., 2017; Nott et al., 2015; Pizzorusso et al., 2002; Stephany et al., 2014). Circuit integration of PV^+^ interneurons, driven by both feed-forward thalamocortical and local intracortical excitation, is critical for the initiation of ODP, while multilevel modulation of their function by different molecular brakes controls critical period closure (Gu et al., 2013; Kuhlman et al., 2013; Takesian and Hensch, 2013; Trachtenberg, 2015). Cell-autonomous synaptic factors that control excitatory inputs on cortical PV^+^ interneurons and thereby the onset and duration of ODP remain uncharacterized. Recent studies have highlighted roles of the ECM protein Narp, as well as PV^+^-expressed NogoR and Neuregulin 1/ErbB4 signaling in the control of local, intracortical excitatory inputs onto PV^+^ interneurons during ODP (Gu et al., 2013; Stephany et al., 2016; Sun et al., 2016). However, the mechanisms that organize feed-forward thalamic inputs onto PV^+^ interneurons and their role in ODP are unknown.

Synapse-organizing adhesion molecules are promising candidates to instruct the assembly of TC excitatory synapses on PV^+^ interneurons. Trans-synaptic interactions between the adhesive pairs of neuroligins and neurexins that can include Hevin as a bridging molecule, cadherins, immunoglobulin proteins of the family of Synaptic Cell Adhesion Molecules (SynCAMs), as well as select other adhesion proteins organize synapses and orchestrate their assembly, maturation, and plasticity across different cell types and brain regions (Missler et al., 2012; Schreiner et al., 2017; Shen and Scheiffele, 2010). Out of this large group of molecules, only SynCAM transcripts were reported to exhibit visual activity-dependent expression in V1 (Lyckman et al., 2008). SynCAMs 1-4 are expressed throughout the CNS, and SynCAM 1 functions in the hippocampus to assemble and maintain synapses on both principal cells and PV^+^ interneurons, and regulates long-term plasticity (Frei et al., 2014; Park et al., 2016; Robbins et al., 2010; Thomas et al., 2008). A role for trans-synaptic organizers in the maturation of cortical inhibition and control of ODP has not been previously described.

Here, we identify SynCAM 1 as the first cell-autonomous synaptic organizer for the development of feed-forward thalamocortical inputs onto PV^+^ interneurons in V1. In this role, SynCAM 1 is essential for PV^+^ interneuron integration and maturation of the TC circuit for vision. Specifically, mice lacking SynCAM 1 exhibit immature inhibitory responses to vision and an ODP that extends beyond the critical period into adulthood. Remarkably, PV^+^-specific knockdown of SynCAM 1 in V1 is sufficient to restore cortical inhibition to earlier developmental levels and extends the critical period. This demonstrates a specific synaptic locus for critical period plasticity in the cortex and points to a central role of feed-forward TC inputs to PV^+^ interneurons in cortical plasticity (Kuhlman et al., 2013). Together, our study determines a PV^+^ cell-autonomous and input-specific mechanism that controls the maturation of the visual circuit and restricts cortical plasticity in the developing and mature brain.

## Results

### Visual activity selectively regulates SynCAM 1 expression levels in V1

SynCAMs are synaptogenic proteins of the immunoglobulin superfamily that form trans-synaptic complexes (Biederer et al., 2002; Fogel et al., 2007; Robbins et al., 2010) and all four SynCAM family members are expressed throughout the brain (Thomas et al., 2008). Their developmental and regional profiles differ, though, and only SynCAM 1 transcripts exhibit an increase in cortical expression after P15, when extensive synaptic remodeling begins (De Felipe et al., 1997; Thomas et al., 2008). To address the extent of developmental protein expression changes in V1, we performed quantitative immunoblotting of total homogenates sampled from V1 of mice at four main stages of V1 development: postnatal day 7 (P7)-start of synaptogenesis; P14-eye opening; P28-peak of the critical period for cortical plasticity; and P45-young adult (Figure 1A) (Kuhlman et al., 2013; Smith and Trachtenberg, 2007). SynCAM 1 protein was present in V1 as early as P7, after which its expression increased strongly through P14 and P28 and remained high in adult mice (Figure 1A).

**Figure 1.**
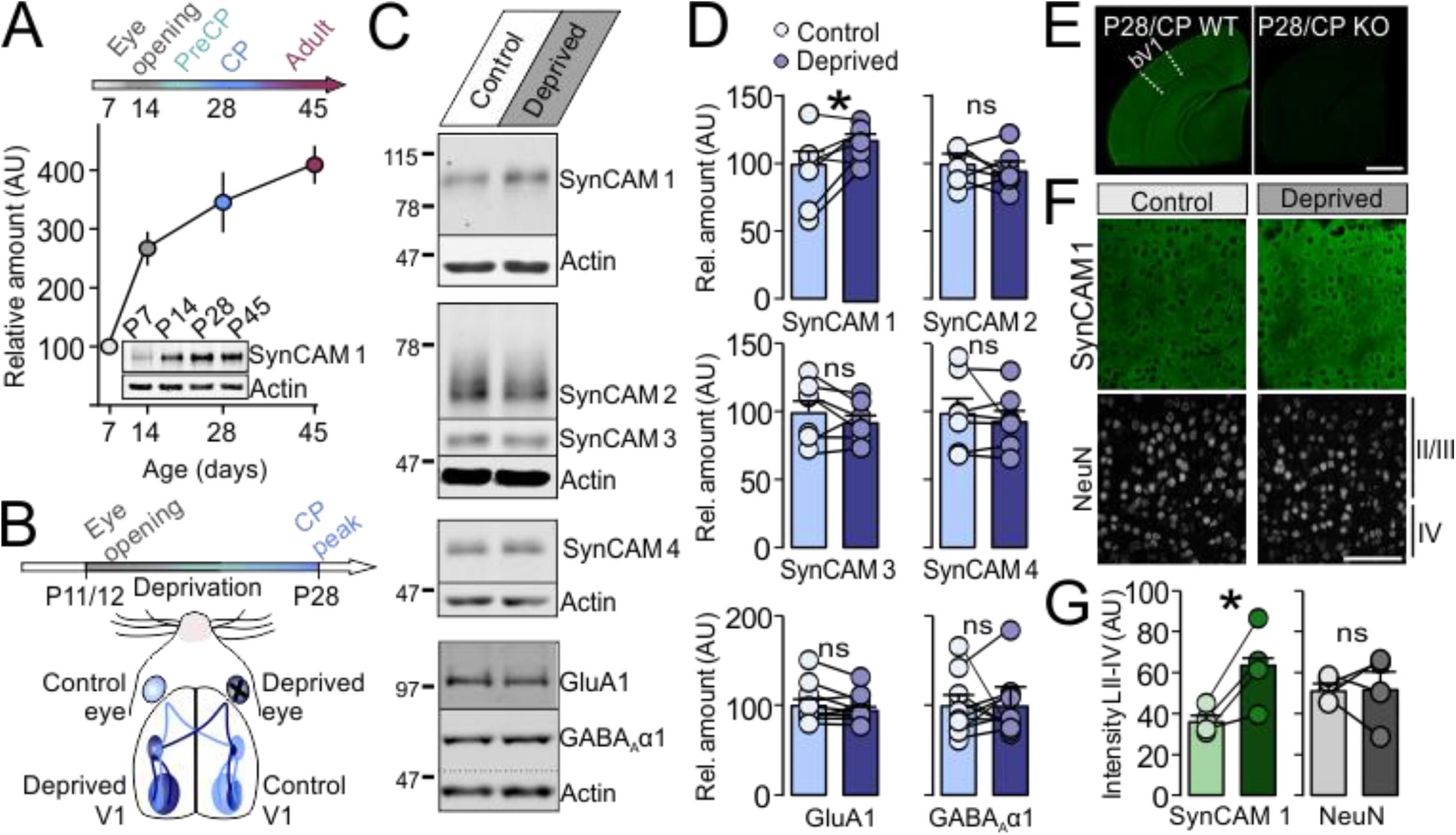
Expression of SynCAM 1 in V1 is regulated by visual experience. A. Quantitative immunoblotting of SynCAM 1 protein expression in the mouse V1 (30 μg/lane). The onset of prominent SynCAM 1 expression coincides with eye opening around P14. SynCAM 1 amounts continue to gradually increase into adulthood (postnatal day 45). N=2 animals/time point. Expression levels were normalized to postnatal day 7 levels. Actin served as loading control. The inset on top marks postnatal stages of cortical development (PreCP, precritical period; CP, critical period).
B. Monocular deprivation (MD) paradigm. Mouse pups had one eye sutured shut at P11/12, before eye opening. Due to the decussation of retinal axons at the chiasm, neurons in the left visual cortex (contralateral to the deprived eye) lose visual responsiveness. The right visual cortex (ipsilateral) continues to receive input from the open eye and served as control in panels C-G.
C. Visual cortices were collected from the control and deprived V1 hemispheres at P28 (critical period/CP peak) as described in B. Quantitative immunoblots of total V1 protein homogenates (30 μg/lane) are shown after probing for SynCAMs 1-4 and GluAl and GABA_A_αl receptors. Molecular weights are indicated on the left. Broad SynCAM bands are due to N-glycosylation (Fogel et al., 2007). Actin served as loading control.
D. SynCAM 1 protein expression is regulated by visual experience. Quantification of data as in (C) showed that monocular deprivation (MD) significantly upregulated SynCAM 1 levels in V1 (top left graph), but had no effect on SynCAMs 2-4 (top right and middle) or on GluA1 and GABA_A_αR1 (bottom panels). Measurements were first normalized to actin and then to control (ipsilateral side) levels. Pairwise expression levels in the same animals are indicated. N=7-10 mice/experiment; *, p<0.05, paired t-test.
E. SynCAM 1 was detected by immunohistochemistry throughout the cortex during the critical period (P28) (left image; binocular V1 delineated). Lack of staining in SynCAM 1 KO animals demonstrated the specificity of the antibody (right). Scale bar: 1 mm
F. Monocular deprivation upregulated SynCAM 1 in layers II-IV (green). The neuronal nuclei marker NeuN served as cell count control (grey).
G. Quantification of data as in (F). N=4 mice/experiment; *, p<0.05, paired t-test. Scale bar: 100 μm. All data presented as mean±SEM, Ns are indicated.

A previous study that sought to identify molecular regulators of critical period plasticity in V1 reported that monocular deprivation by eyelid suture throughout the critical period intensely upregulated SynCAM gene expression in V1, with SynCAMs the second most strongly regulated transcripts (Lyckman et al., 2008). To determine which of the SynCAMs is regulated by visual activity on the protein level, we performed quantitative immunoblotting of V1 tissue samples in mice that had undergone monocular deprivation from the beginning of eye opening at P11/12 until the peak of the critical period (Figure 1B) (Lyckman et al., 2008; Tropea et al., 2006). V1 of mice is strongly driven by contralateral eye, with a small binocular region containing neurons responsive to both eyes (Figure 1B). This connectivity pattern allows for intra-animal comparison of changes in protein expression, where V1 contralateral to the deprived eye is impacted by deprivation, while the ipsilateral cortex serves as non-deprived control (Figure 1B) (Heynen et al., 2003). Only SynCAM 1 exhibited a significant activity-dependent change in expression (Figure 1C and 1D). Monocular deprivation upregulated SynCAM 1 protein levels (Figure 1C and 1D; Control V1=100±11 arbitrary units (AU), Deprived V1=120±5.3 AU, paired t-test p=0.042; N=7 animals, t=2.6, df=6), but had no effect on SynCAM 2, 3 and 4 (Figure 1C and 1D; SynCAM 2: Control=100±6.8, Deprived=95±7.2; SynCAM 3: Control=100±8.2, Deprived=92±5.1; SynCAM 4: Control=100±10, Deprived=94±7.7; all values in AU). Consistent with previous reports, deprivation did not affect the levels of glutamate or GABA receptors (Figure 1C and 1D; GluA1: Control=100±6.9, Deprived=95±4.8; GABA_A_αl: Control=100±11, Deprived=100±11; paired t-test, N=10 animals; all values in AU) (Lyckman et al., 2008; Tropea et al., 2006). SynCAM 1 therefore shows a selective expression profile in V1 that is regulated during development and modulated by visual activity.

We used SynCAM 1-specific antibodies suitable for immunohistochemistry to evaluate activity-dependent changes in protein expression (Figure 1E). Quantitative immunohistochemistry of sections through V1 from monocularly deprived mice confirmed that deprivation strongly upregulated SynCAM 1 expression in thalamorecipient layers II/III and IV, while the neuronal marker NeuN remained unaffected (Figure 1F and 1G; SynCAM 1 Control=37±3.2 AU, Deprived=63±9.7 AU, paired t-test p=0.033; N=4 animals, t=3.8 df=3; NeuN Control=52±2.8 AU, Deprived=52±8.8 AU). These results are consistent with the changes detected using immunoblotting and demonstrate that SynCAM 1 protein expression is regulated by visual activity during the critical period in V1.

### SynCAM 1 limits visual plasticity in both juvenile and adult brain

Monocular deprivation by eyelid suture during the critical period robustly depresses closed (contralateral) eye responses after 3-4 days, resulting in a strong downward shift in the contra/ipsi (C/I; closed/open) eye response ratio (Frenkel and Bear, 2004; Gordon and Stryker, 1996). Studies indicate that long-term depression (LTD) mediates the decrease in closed eye response to deprivation (Frenkel and Bear, 2004; Rittenhouse et al., 1999). SynCAM 1 restricts hippocampal LTD (Robbins et al., 2010) and visual activity strongly regulated SynCAM 1 expression (Figure 1D and 1G). We therefore hypothesized that SynCAM 1 may modulate deprivation-induced plasticity in V1. To test this, we recorded visually evoked potentials (VEPs) from the binocular V1 in awake, freely-behaving animals using a spherical treadmill setup (Figure 2A and 2B) (Dombeck et al., 2007; Niell and Stryker, 2010), thus abrogating the potential effects of anesthetics on cortical plasticity (Hubener and Bonhoeffer, 2014). We monocularly deprived SynCAM 1 knock-out (KO) animals and their wild type (WT) littermates by eyelid suturing for 3-4 days before the onset of the critical period (P21-24, PreCP), at the peak of the critical period (P25-28, CP) and in adult animals (P60-64; Figure 2C). We reopened the sutured eyelid on the last day of deprivation and recorded visual responses to the stimulation of both closed (contra) and open (ipsi) eyes to calculate the C/I ratio. Naïve, non-deprived WT and KO animals had almost identical C/I ratios (Figure 2D, WT=2.4±0.2, KO=2.6±0.3, Supplementary Table 1) and similar VEP amplitudes (Figure 2E, WT Contra /right eye=183±11.4 μV, KO Contra /right eye=169±16.6 μV; WT Ipsi/left eye=78±6.4 μV and KO Ipsi/left eye=69±6.9 μV; Supplementary Table 1), suggesting that visual responses are grossly normal in the absence of SynCAM 1. Consistent with previous studies, 3 days of deprivation during the precritical period were not sufficient to significantly affect visual responses in WT animals (Figure 2D and 2E, C/I PreCP=1.8±0.3; Contra/closed eye=145±27.3 μV, Ipsi/open eye=86 ±17.3 μV, Supplementary Table 1) (Hanover et al., 1999). Deprivation of WT mice during the critical period induced the expected robust shift in C/I ratio and strong depression of closed eye responses (Figure 2D and 2E, MD CP C/I=1.5±0.2, p=0.022; Contra/closed eye=115±21.6 μV, p=0.05; Ipsi/open eye=72±11 μV, p=ns; Supplementary Table 1) (Frenkel and Bear, 2004; Gordon and Stryker, 1996). Short-term deprivation had no effect in adult WT animals, in agreement with the reduced plasticity of the mature cortex (Figure 2D and 2E, MD Adult C/I=2.1±0.2, Contra/closed eye=177±25.9 μV, Ipsi/open eye=89±16.7 μV; Supplementary Table 1) (Lehmann and Lowell, 2008; Morishita et al., 2010).

**Figure 2.**
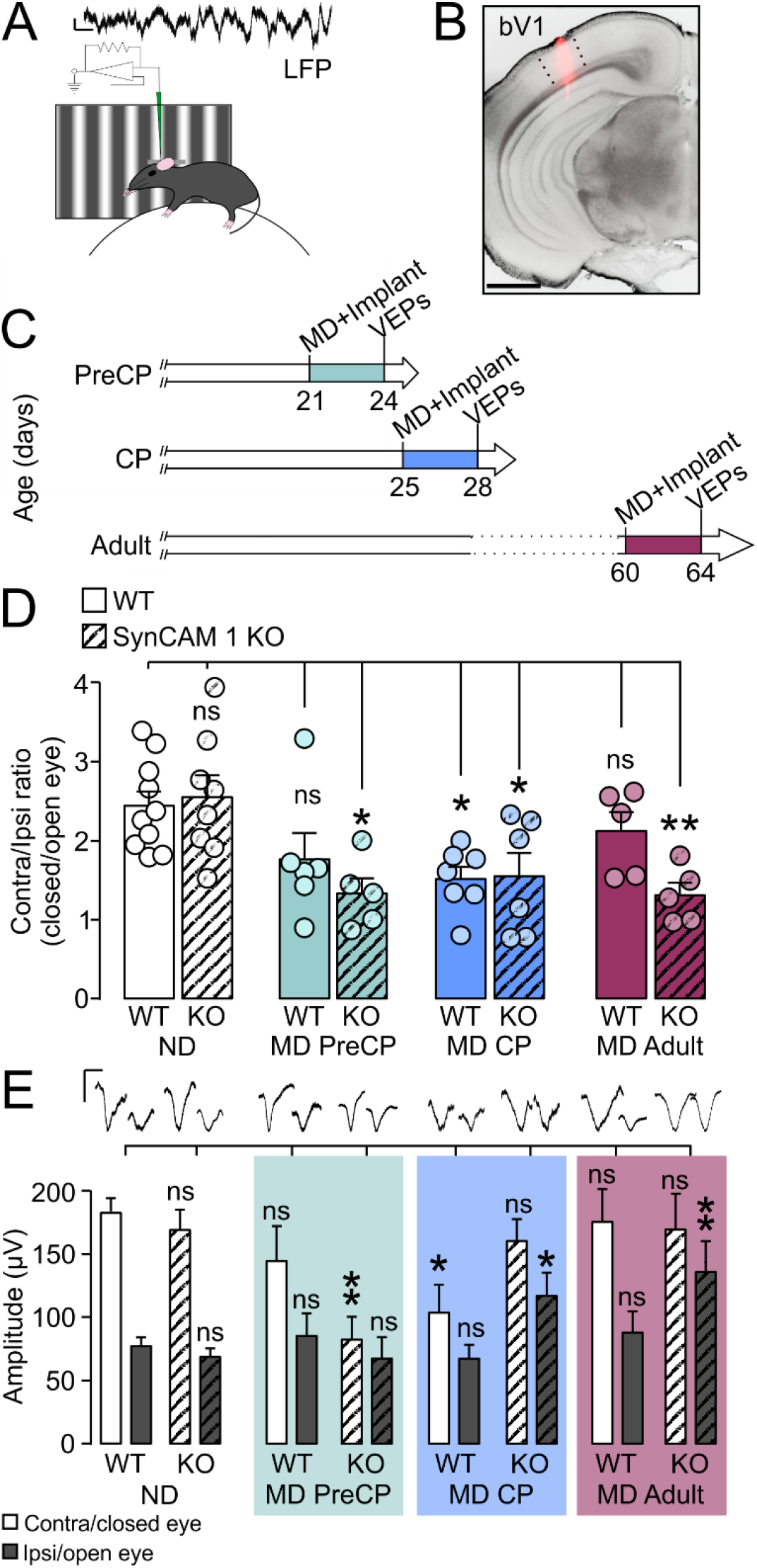
SynCAM 1 limits plasticity in the visual cortex. A. During *in vivo* VEP recordings using spherical treadmill, animals are head-fixed over an air-suspended styrofoam ball. Full field visual stimuli are presented and visually evoked local field potentials (LFPs) are simultaneously recorded using 16-channel probes. Scale bars: 250 μV and 0.5 s.
B. Electrode tracts were labeled with DiI (pink) and overlaid on DAPI signal (greyscale inverted). The image shows a representative electrode tract with penetration through all cortical layers in the binocular V1. Scale bar: 1 mm.
C. Timelines of MD in this study. Mice at the indicated postnatal days (PreCP at P21, CP at P25, and adult at P60) had their right eyes sutured shut and titanium headposts for *in vivo* physiology were attached to the skull. The deprived eye was reopened 3-4 days later, prior to the recording of VEPs. Non-deprived animals underwent parallel surgical procedures without eyelid suture.
D. Binocular V1 is contralaterally driven, reflected in a high contralateral/ipsilateral (C/I) VEP amplitude ratio in non-deprived (ND) WT mice at P28 (white bars). C/I ratio of non-deprived SynCAM 1 KO mice was comparable to WT mice at P28, indicating normal gross visual function (white/striped bars). MD of WT mice during the PreCP resulted in a trend towards lower C/I ratio that was not significant (green), while PreCP SynCAM 1 KO mice displayed a robust C/I reduction after deprivation (green/striped), indicating heightened plasticity. Both WT and SynCAM 1 KO mice were plastic during the CP, when MD caused a significant reduction in the C/I ratio (blue and blue/striped). Plasticity in WT mice tapered in adulthood when short MD had no effect on visual responses (crimson). In contrast, C/I of adult SynCAM 1 KO was significantly lowered after monocular deprivation (crimson/striped), indicating an extended critical period.
E. VEP amplitudes elicited by the stimulation of deprived (contra) and open (ipsi) eyes in the left binocular V1 after right eye suture. Data were obtained from recordings as in D. VEP amplitudes of WT mice (white bars) were similar to non-deprived SynCAM 1 KO mice (striped bars), indicating normal gross visual function in the KO. MD during the PreCP (green background) in WT mice had no significant effect on the closed eye VEP (white bars), but caused strong depression of the closed eye responses in KO mice (striped bars). MD during the CP (blue background) significantly depressed the closed eye responses in WT mice, but increased the open eye responses in KO mice (striped bars). MD in adult (crimson background) WT mice had no effect (empty bars), while significantly potentiating open eye responses in SynCAM 1 KO mice (striped bars). Scale bars: 100 μV and 0.2 s. Panels D, E: ns, not significant; *, p<0.05; * *, p<0.01; One-way and Two-way ANOVA (see Supplementary Table 1 for details). Data presented as mean±SEM, Ns are indicated.

Unlike WT mice, short monocular deprivation decreased C/I ratio at all ages tested in SynCAM 1 KO mice (Figure 2D and 2E, MD PreCP=1.3±0.2, p=0.011; MD CP=1.6±0.3, p=0.039; MD Adult=1.3±0.16, p=0.009; Supplementary Table 1). 3 days of deprivation in SynCAM 1 KO mice strongly depressed closed eye responses even during the precritical period, while inducing open eye potentiation during the critical period (Figure 2E, PreCP Contra/closed eye=83±18.2 μV, p=0.005; PreCP Ipsi/open eye=68±17.1 μV, p=ns; CP Contra/closed eye=161±17.5 μV, p=ns; CP Ipsi/open eye=118±18.3 μV, p=0.033; Supplementary Table 1). In striking contrast to WT mice, both critical period and adult SynCAM 1 KO mice had plastic responses to monocular deprivation, and these were almost identical with significant open eye potentiation in both age groups (Figure 2E, Adult Contra/closed eye=170±27.7 μV, Adult Ipsi/open eye=136±23.9 μV, p=0.004; Supplementary Table 1). Two-way ANOVA analysis demonstrated a significant interaction between genotype and deprivation in the amplitude of open (ipsi) eye responses (F_(3,44)_=3.1, p=0.035). Short deprivation at P17 had no effect on either WT or KO mice (Supplementary Figure 1), suggesting that the critical period was not shifted to an earlier time in KO mice (Hanover et al., 1999; Miyata et al., 2012). These data specifically implicate SynCAM 1 is restricting the closure, but not the opening of the critical period.

### Formation of perineuronal nets is impaired in the absence of SynCAM 1

The closure of the critical period in V1 requires the maturation of PV^+^-mediated cortical inhibition (Fagiolini et al., 2004; Kuhlman et al., 2013). As PV^+^ interneurons mature, proteoglycan-composed extracellular matrix (ECM) structures called perineuronal nets (PNNs) form around them (Figure 3A) (Lensjo et al., 2017; Pizzorusso et al., 2002; Sorg et al., 2016). Adult SynCAM 1 KO mice retained the potential for robust plasticity even after critical period closure, similar to what has been observed after removal of the ECM (Pizzorusso et al., 2002) (Figure 2D and 2E). We studied the development of PNNs in SynCAM 1 KO V1 by quantifying the staining intensity of *Wisteria floribunda* agglutinin (WFA), a marker for PNNs (Lensjo et al., 2017; Pizzorusso et al., 2002; Sugiyama et al., 2008). At the start of the precritical period (PreCP), WT mice already had over 60% of their PV^+^ interneurons enwrapped with PNNs (Figure 3B and 3D, WT PreCP/P21=67±6.3, N=5 animals; WT CP/P28=70±4.5, N=4; WT Adult/P60-70=74±2, N=4; all values in % of PV^+^ interneurons). In addition, the density of those PNN puncta around PV^+^ interneurons that were positive for PNNs steeply increased from precritical period to adulthood in WT mice (Figure 3E; WT PreCP/P21=301±61.7, CP/P28=837±139.3, Adult/P60-70=1271± 113.2; all values in particles/mm^2^). The overall density of PV^+^ interneurons in SynCAM 1 KO mice was indistinguishable from WT mice (Figure 3F; PreCP P21 WT=141±7, KO=140±14.1; CP/P28 WT=163±10.2, KO=168±15.5; Adult/P60-70 WT=155±4.8, KO=155±8.9; all values in cells/mm^2^). In contrast to the prominent enwrapping of PV^+^ cells in WT mice, the fraction of PV^+^ interneurons surrounded by PNNs was significantly lower in SynCAM 1 KO mice at all ages studied (Figure 3D; PreCP/P21=42±5.5, p=0.008; CP/P28=35±3.7, p=0.0006; Adult/P60-70=50±5, p=0.014; One-way ANOVA with Holm-Sidak’s multiple comparisons test F_(5,16)_=9.9, p=0.0002). In addition, the loss of SynCAM 1 severely reduced PNN deposition that started to be significant at the peak of the critical period (P28) (Figure 3E, KO PreCP/P21=75±47.5, N=3, p=0.415; KO CP/P28=178±50.4, N=3, p=0.003; KO Adult/P60-70=766±195.2, N=3, p=0.019; all values in particles/mm^2^; One-way ANOVA with Holm-Sidak’s multiple comparisons test F(5,16)=17.4, p<0.0001). Development of PNNs is hence impaired in the absence of SynCAM 1 from the precritical period through adulthood.

**Figure 3.**
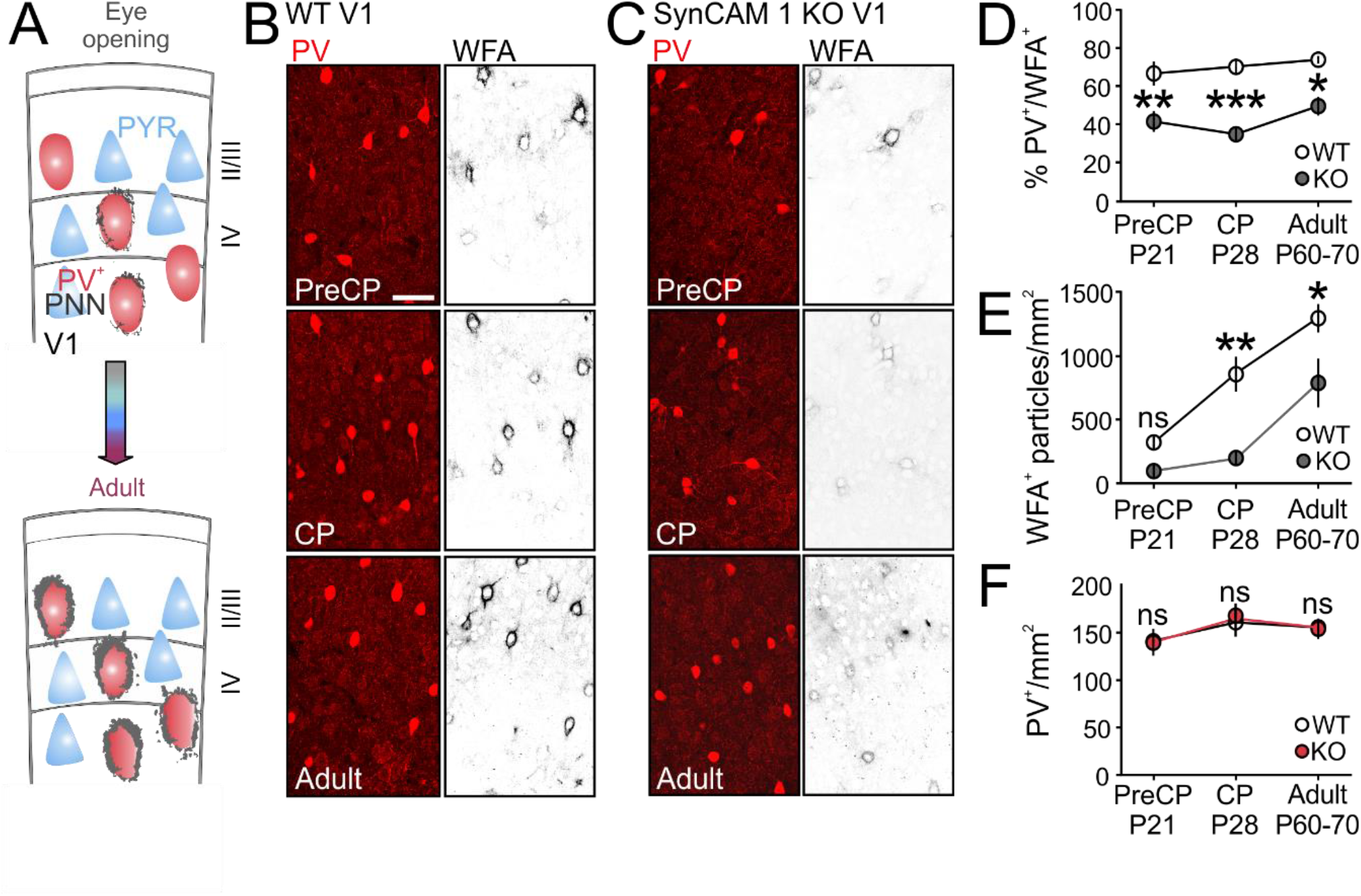
Formation ofPNNs is reduced in the absence of SynCAM 1. A. The model illustrates that prior to eye opening, few PV^+^ interneurons (red) are surrounded by PNNs (grey). After eye opening, PNN deposition increases parallel to the maturation of PV^+^ interneurons. Pyramidal neurons (PYR; blue) lack PNNs.
B. Comparison of single optical confocal sections after double-labeling for PV (red) and WFA (gray), a marker of PNNs, in WT V1 at the indicated ages (PreCP at P21, CP at P28 and Adult at P60-70) showed that WFA^+^ PNNs enwrapped the majority of PV^+^ interneurons in the WT. Scale bar: 50 μm.
C. PNNs surrounded fewer PV^+^ interneurons in SynCAM 1 KO mice at all ages tested as imaged in single optical confocal sections. The density of PNN particles around those PV^+^ cells that were positive for WFA was also lower in KO animals.
D. Quantification of images as in (B) and (C) showed a significant difference in the density of WFA-positive PV^+^ interneurons between genotypes through all ages tested.
E. The density of WFA^+^ particles that comprise the PNNs increased throughout age in WT animals, but remained significantly lower from the critical period onwards in the fraction of PV^+^ cells of SynCAM 1 KO mice that remained positive for WFA.
F. The density and distribution of PV^+^ cells were indistinguishable between WT and SynCAM 1 KO. Panels D-F: ns, not significant; *, p<0.05; **, p<0.01; ***, p<0.001; One-way ANOVA. Data presented as mean±SEM, N=3-5 animals/genotype.

PNN density in SynCAM 1 KO mice was lowered at all stages and even in the adult did not reach WT critical period levels (Figure 3E). To assess whether SynCAM 1 is a molecular component of PNNs, we analyzed WFA and SynCAM 1 staining in cortical cultures. We observed strong SynCAM 1 staining in both immature WFA^-^ PV^+^ interneurons as well as WFA^+^ PV^+^ interneurons in mature cultures that overlapped but did not selectively colocalize with WFA (Supplementary Figure 2). This indicated that SynCAM 1 is not a component specific to PNNs.

A PNN-dependent transcriptional trigger for the maturation of PV^+^ interneurons is the non-cell autonomous homeobox transcription factor Otx2 (Beurdeley et al., 2012; Sugiyama et al., 2008). Otx2 can be acquired by PV^+^ interneurons through PNNs to create a positive feedback loop between internalized Otx2 and PNN deposition (Sugiyama et al., 2008). The density of PV^+^ interneurons positive for Otx2 was indistinguishable between WT and SynCAM 1 KO mice, indicating that altered Otx2 expression does not account for the decreased PNN formation in SynCAM 1 KO mice (Supplementary Figure 3).

### SynCAM 1 is necessary for recruitment of thalamocortical terminals onto PV^+^ interneurons

Neuronal activity controls the deposition of PNNs (Dityatev et al., 2007). Specifically, the inhibition of fast, α-amino-3-hydroxy-5-methyl-4-isoxazolepropionic acid (AMPA)-receptor mediated excitatory synaptic transmission *in vitro* (Dityatev et al., 2007), as well as the blockade of visual input by dark rearing *in vivo*, reduce the density of PNNs around PV^+^ interneurons (Ye and Miao, 2013). Given reduced PNN formation in the absence of SynCAM 1 and the roles of SynCAM 1 in the development of excitatory inputs on both pyramidal cells and interneurons in the hippocampus (Park et al., 2016; Robbins et al., 2010), we investigated glutamatergic synaptic inputs on cortical PV^+^ interneurons in SynCAM 1 KO mice. SynCAM 1 was intensely expressed in PV^+^ interneurons both *in vitro* (Supplementary Figure 2) and *in vivo* (Figure 4A), where it formed dense dendritic puncta. PV^+^ interneurons in the cortex receive two classes of excitatory synapses, short-range, intracortical and long-range, thalamocortical (TC) (Figure 4B) (Fremeau et al., 2001). Intracortical synapses utilize the presynaptic vesicular glutamate transporter 1 (vGlut1), whereas long-range TC inputs are marked by vGlut2 (Figure 4B) (Coleman et al., 2010; Fremeau et al., 2004; Kameda et al., 2012; Singh et al., 2016). Confocal microscopy enables reliable measurements of the density of vGlut1 *vs* vGlut2 puncta in contact with PV^+^ dendrites (Kameda et al., 2012; Park et al., 2016; Pizzo et al., 2016; Sonomura et al., 2013). WT mice exhibited intense vGlut1 signal throughout the cortex, while vGlut2 was most intense and appeared as a thick band in the thalamorecipient layer IV, consistent with previous studies (data not shown) (Coleman et al., 2010; Fremeau et al., 2004). The number of vGlut1 and vGlut2 puncta in contact with PV^+^ dendrites in layer IV of WT mice did not vary from precritical period (PreCP/P21) to adulthood (P60-70) in agreement with previous reports (Figure 4C and 4D; PreCP/P21=18±0.4, N=5 animals; CP/P28=17±0.5, N=4; Adult/P60-70=17±0.7, N=4; all values in puncta/100 μm^2^) (Kameda et al.,2012). Importantly, the lack of SynCAM 1 in KO mice significantly reduced vGlut2^+^ TC inputs to PV^+^ interneurons from the precritical period onwards compared to WT littermate controls, with this difference being most pronounced in adult animals (Figure 4C) (KO PreCP/P21=16±0.6, N=3, p=0.036, 12% reduction; CP/P28=14±0.5, N=3, p=0.008, 19% reduction; Adult/P60-70=13±1.3, N=3, p=0.001, 25% reduction; all values in puncta/100 μm^2^; One-way ANOVA with Holm-Sidak’s multiple comparisons test F(5,16)=8.5, p=0.0004). In contrast, the density of intracortical, vGlut1^+^ inputs was indistinguishable between WT and KO animals at all ages (Figure 4D) (PreCP/P21 WT=35±0.8, KO=33±0.9; CP/P28 WT=31±1.2, KO=30±1.8; Adult/P60-70 WT=33±1.7, KO=34±0.4; all values in puncta/100 μm^2^). Puncta size was not significantly different between the groups for both vGlut1 and vGlut2 (data not shown). These results suggest a TC input-specific abnormality of PV^+^ interneurons in the absence of SynCAM 1.

**Figure 4.**
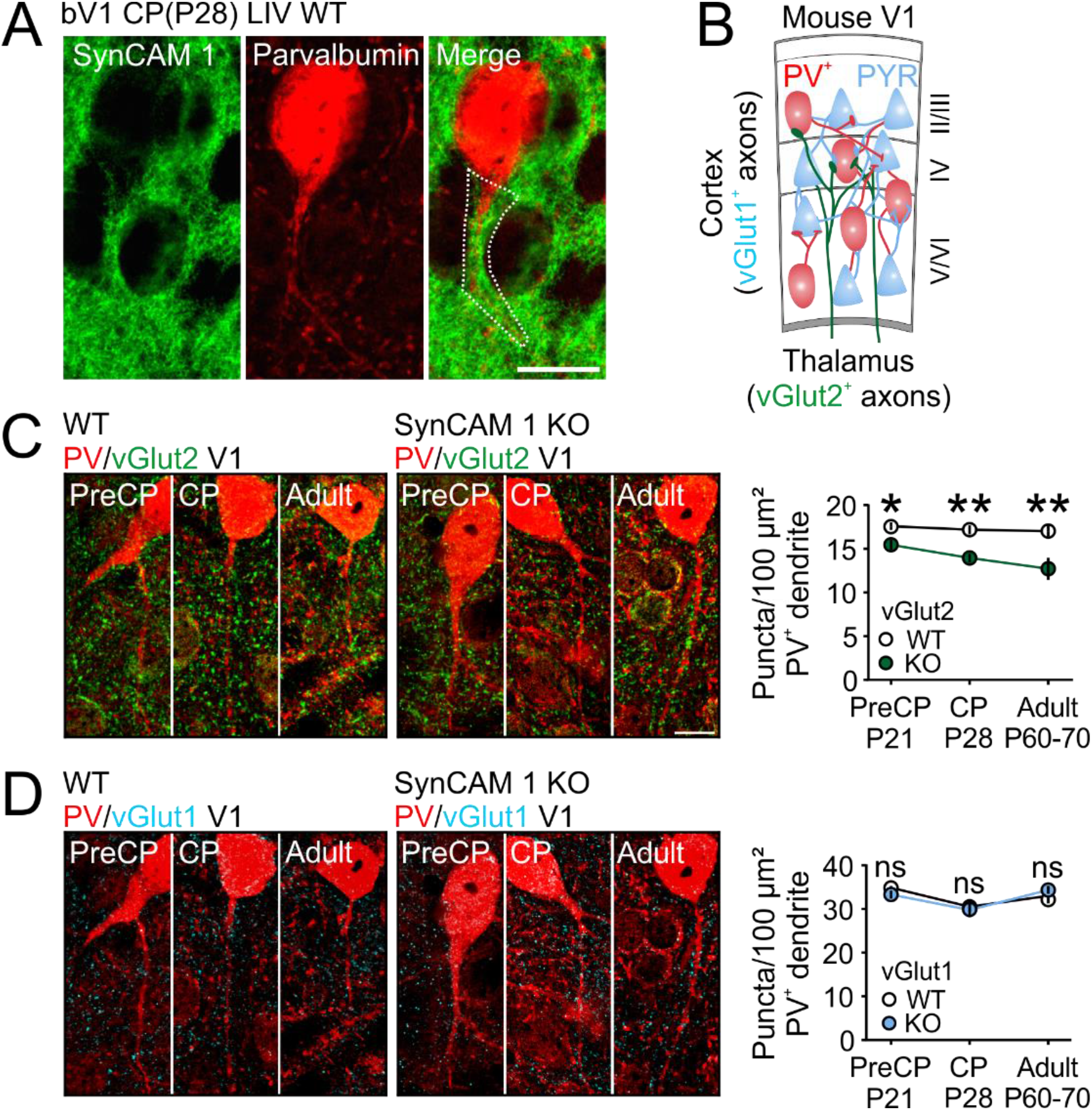
PV^+^ interneurons in V1 of SynCAM 1 KO mice receive fewer inputs from thalamus. A. SynCAM 1 (green) is abundant in the neuropil in the critical period cortex, where it localizes to dendrites of PV^+^ interneurons (red). Images represent a single optical section. Scale bar: 15 μm.
B. Connectivity of mouse V1. vGlut2^+^ inputs from dLGN (green) terminate mainly in layers II/III and IV, where they innervate both PV^+^ inhibitory neurons (red) and pyramidal neurons (blue). Local cortico-cortical connections predominantly use vGlut1 (cyan).
C. Representative single optical sections of PV (red)/vGlut2 (green) immunofluorescence in layer IV bV1 of WT and SynCAM 1 KO mice at the indicated ages (PreCP at P21, CP at P28 and Adult at P60-70). The graph on the right depicts the density of vGlut2^+^ puncta in contact with PV^+^ dendrites as measured from images as on the left. SynCAM 1 KO mice showed a significant reduction in TC inputs in contact with PV^+^ dendrites at all ages. Scale bar: 5 μm.
D. Representative single optical sections of PV (red)/vGlut1 (green) immunofluorescence in layer IV bV1 of WT and SynCAM 1 KO mice at the indicated ages (PreCP at P21, CP at P28 and Adult at P60-70), imaged and analyzed asin (C). SynCAM 1 KO and WT PV^+^ cells had an indistinguishable density of intracortical vQut1^+^ inputs. Panels C, D: ns, not significant; *, p<0.05; **, p<0.01; ***, p<0.001; One-way ANOVA. Data presented as mean±SEM, N=3-5 animals/genotype.

### The maturation of the visual circuit involves SynCAM 1

Synaptic transmission at TC synapses activates PV^+^ interneurons more intensely than pyramidal neurons, giving rise to strong feed-forward inhibition in the cortex (Cruikshank et al., 2007; Ji et al., 2016; Kloc and Maffei, 2014). This sensory activity-driven transmission at TC synapses is critical for cortical circuit maturation (Chittajallu and Isaac, 2010). In V1, feed-forward inhibition matures after eye opening and strongly suppresses the primary response to visual stimulation (Shen and Colonnese, 2016). This effect of feed-forward inhibition is often inconspicuous in VEP measurements, but becomes evident upon analysis of action potential firing rates at the level of both single and multiple neurons (Gu et al., 2013; Shen and Colonnese, 2016). To determine how the reduced density of TC inputs onto PV^+^ interneuron synapses in the absence of SynCAM 1 affects the maturation of visual responses, we analyzed spontaneous and stimulus-induced activity of neurons (multi-unit activity (MUA)) in layer IV from filtered VEPs (Figure 5A). During the critical period, both WT and KO animals showed robust and transient increases in firing rate in response to visual stimulation (Figure 5B and 5C) (Shen and Colonnese, 2016; Gu et al., 2013). Spontaneous firing rate was indistinguishable between SynCAM 1 KO and WT (Figure 5A-C and 5D; spontaneous (average prestimulus) firing: WT=15±1.7 Hz, KO=15±1.5 Hz, Supplementary Table 2). However, the stimulus-evoked firing rate was significantly increased in SynCAM 1 KO animals (Figure 5A-C and 5E; evoked firing rate=peak poststimulus-spontaneous firing rate; WT=21±2.7 Hz, KO=34±4.4 Hz, p=0.002; Supplementary Table 2), indicating disinhibition of visual responses (Gu et al., 2013; Kaplan et al., 2016). Detailed analysis of firing revealed a significantly protracted primary visual response in SynCAM 1 KO animals (Figure 5F-H; primary response latency: WT=23±2.3 ms, KO=35±3.3 ms, p=0.004; primary response duration: WT=23±3.2 ms, KO=43±4.8 ms, p<0.001; Supplementary Table 2). Some feed-forward inhibition was still evident in SynCAM 1 KO V1 (Figure 5C and 5H), consistent with persistence of a fraction of TC inputs onto PV^+^ interneurons (Figure 4C), but the effect of inhibition was significantly delayed (Figure 5C and 5H; latency of feedforward inhibition: WT=46±4.8, KO=78±6.3, p<0.001; Supplementary Table 2). Feed-forward inhibition in V1 is therefore impaired when SynCAM 1 is absent, consistent with the reduction in feed-forward TC excitation onto PV^+^ interneurons.

**Figure 5.**
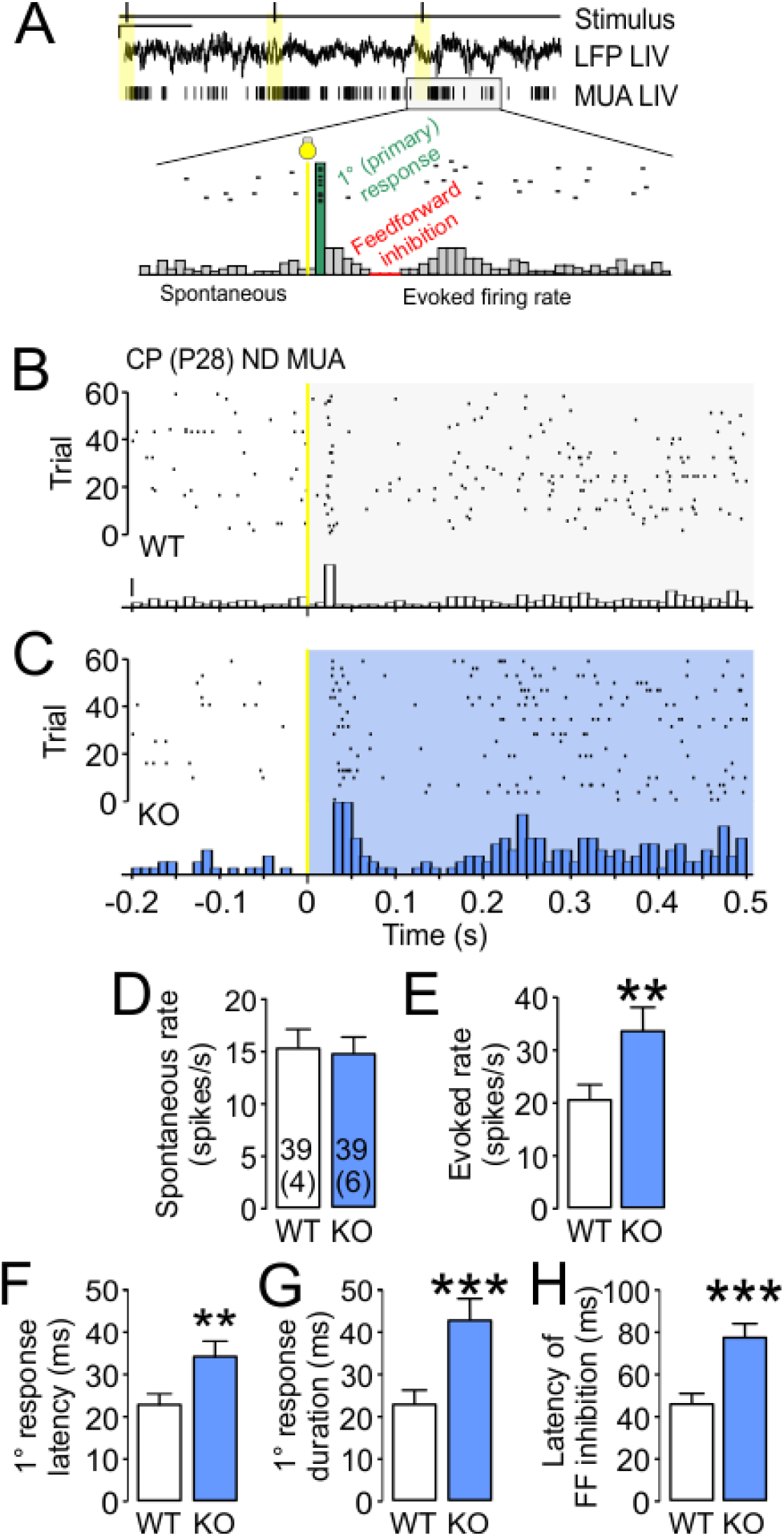
Feed-forward inhibition in V1 is immature in SynCAM 1 KO mice. A)MUA was extracted from filtered VEPs from layer IV. Stimulus (LED flash) is indicated in yellow. The enlarged inset shows an example of spiking activity and peristimulus time histogram (PSTH), and marks measured parameters. Evoked firing rate was calculated as average spontaneous firing rate subtracted from average peak firing rate. Scale bars: 100 μV and 2.5 s.
B,C) Representative raster plots and PSTHs of MUA recorded from WT (B) and SynCAM 1 KO mice (C) at P28, the peak of CP. Stimulus onset is indicated in yellow. Late onset of firing and prolonged primary response was evident in SynCAM 1 KO animals. Scale bar: 15 spikes/s.
D-E) Average spontaneous, prestimulus firing rate was comparable in SynCAM 1 KO mice to that of WT mice (D), but the evoked firing rate was significantly increased in SynCAM 1 KO animals (E).
F-G) Latency of the primary response (F) and response duration (G) were increased in SynCAM 1 KO mice.
H)Inhibition of the primary response occurred later in SynCAM 1 KO than in WT animals. Panels D-H: **, p<0.01; ***, p<0.001; Mann-Whitney rank sum test; Ns of MUAs and animals/genotype are indicated in D (see Supplementary Table 2 for details). Data presented as mean±SEM.

The maturation of PV^+^-mediated inhibition drives the development of cortical oscillatory activity in the high gamma range (>40 Hz) (Cardin et al., 2009; Chen et al., 2015). In V1, visual stimulation suppresses gamma power during the critical period, which can be prevented by delaying circuit maturation with dark rearing (Chen et al., 2015; Chen et al., 2017). Further, removal of PNNs is correlated with increased plasticity and gamma oscillations (Lensjo et al., 2017). As visual circuits appeared immature in the absence of SynCAM 1 (Figure 5), we hypothesized that gamma range power might be aberrant in SynCAM 1 KO mice during the critical period. Consistent with previous studies (Chen et al., 2015), critical period WT mice exhibited a drop in gamma power (40-70 Hz) after switching their stimulus from blank grey screen to full field sinusoidal gratings (Figure 6A and 6B; WT blank=0.25±0.06 μV^2^, gratings=0.09±0.02 μV^2^, p=0.026; N=11 animals; paired t-test, t=2.6, df=10). Gamma suppression was most pronounced in layer IV (Δpower^blank-ěratings^ at 400 μm=0.21±0.08 μV^2^; Layer II/III at 100-350 μm=0.12±0.06 μV^2^; deep layers V/VI at 450-700 μm=0.13±0.08 μV^2^) (Chen et al., 2015). No change in the low frequency range (1-20 Hz) was measured in WT mice after visual stimulation, as expected (Figure 6A and 6C; WT blank=85±1.6 μV^2^, gratings=85±2.6 μV^2^) (Chen et al., 2015). Similar to WT mice, SynCAM 1 KO mice showed no stimulation-induced changes in the 1-20 Hz band (Figure 6A and 6C; KO blank=80±4.7 μV^2^, gratings=87±1.9 μV^2^, N=8). Interestingly, SynCAM 1 KO mice lacked the visual stimulation-induced suppression of gamma band activity (Figure 6A and 6B; KO blank=0.25±0.07 μV^2^, gratings=0.21±0.06 μV^2^). These results are further evidence that thalamocortical circuitry remains immature in the absence of SynCAM 1.

**Figure 6.**
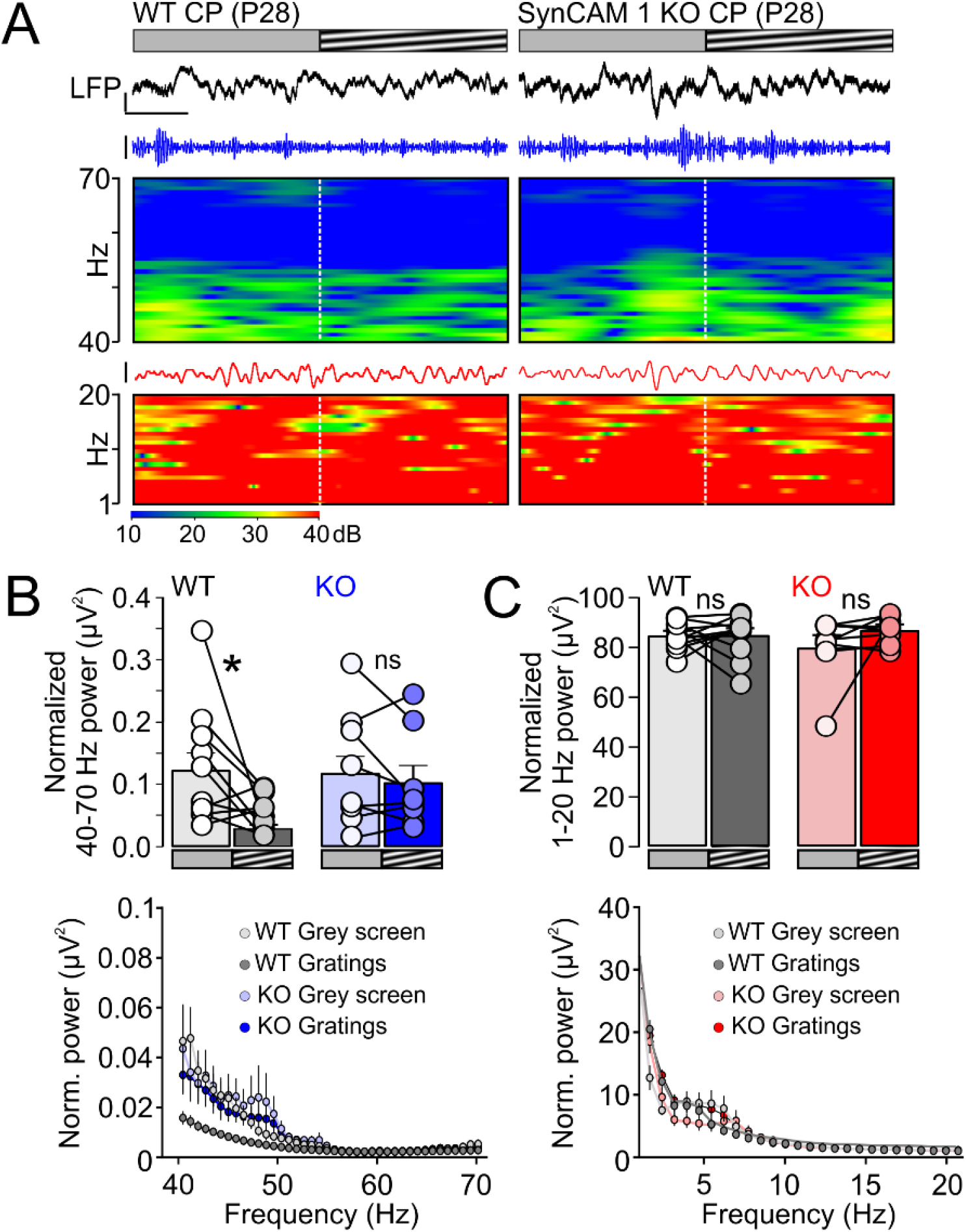
Stimulus-induced suppression ofγ-oscillations during the CP is absent in V1 of SynCAM 1 KO mice. A. Filtering of VEPs uncovers oscillatory network activity. Top, animals were first presented with a grey screen and then shown sinusoidal gratings, and LFPs were recorded. Representative traces were filtered to depict oscillations in γ-range (40-70 Hz, blue) and in lower frequency bands (1-20 Hz; red), as well as sonograms. Suppression of 40-70 Hz oscillations after presentation of sinusoidal gratings was evident in WT sonograms. Scale bars: 350 μV for LFP, 50 μV for 40-70 Hz and 250 μV for 1-20 Hz and 0.5 s.
B. Top: visual stimulation with sinusoidal gratings of varying frequencies robustly suppressed oscillations in the γ-range (40-70 Hz) in WT animals during the CP, compared to grey screen presentation. This effect was absent in SynCAM 1 KO animals. Bottom: histograms of power demonstrated a strong reduction of 40-70 Hz oscillations in the WT after stimulus presentation, but the effect was absent in the KO.
C. Top: no change in lower frequency bands (1-20 Hz) was detected in either WT or SynCAM 1 KO animals after stimulus presentation. Bottom: histograms of power showed no changes in the 1-20 Hz band. Panels B, C: ns, not significant; *, p<0.05; paired t-test. Data presented as mean±SEM, N=11 WTs and 9 KOs.

### SynCAM 1 acts in PV^+^ interneurons in V1 to close the visual critical period

Where does SynCAM 1 function to promote cortical network maturation? We addressed this question through region-and cell-type specific manipulations. Recent studies implicated the visual thalamus in the regulation of critical period plasticity (Jaepel et al; 2017; Sommeijer et al., 2017). SynCAM 1 expression in the lateral geniculate nucleus (LGN) was low and barely detectable during postnatal development (Supplementary Figure 4) and the organization of LGN in SynCAM 1 KO mice was grossly normal (Supplementary Figure 5). These findings suggest a cortical locus for SynCAM 1 action in the regulation of the visual critical period. Cortical fast spiking PV^+^ interneurons in SynCAM 1 KO have fewer synaptic inputs and PNNs (Figures 3 and 4), while regular spiking somatostatin interneurons appeared normal (data not shown), in agreement with high expression of SynCAM 1 in PV^+^ cell population (Foldy et al., 2016). Therefore, we next aimed to directly test whether the locus of SynCAM 1 action in critical period regulation are cortical PV^+^ interneurons. To address this, we cloned a shRNA against SynCAM 1 (Cadm15) (Faraji et al., 2012) and a control scrambled sequence into an adenoviral vector that allows Cre-mediated activation of shRNA expression (Supplementary Figure 6A) (Wohleb et al., 2016). The efficiency of SynCAM 1 knockdown (KD) was validated in heterologous HEK293 cells and in cultured neurons (Supplementary Figure 6B and 6C). To analyze the roles of SynCAM 1 in PV^+^ interneurons, we injected the AAV-shCadm15 into the left cortices of PV-Cre mice (Figure 7A and 7B), where Cre recombinase is driven by the PV^+^ promoter (Hippenmeyer et al., 2005). We injected the KD viral constructs at P14 (Figure 7A), when SynCAM 1 expression normally increases in V1 (Figure 1A). Quantitative immunohistochemistry confirmed Cre-mediated knockdown of SynCAM 1 in adult PV-Cre mice (Supplementary Figure 6D). We then deprived the right (contra) eyes of adult P60 PV-Cre mice and recorded VEPs in awake, freely behaving animals after reopening the right eye 4 days later, as in adult SynCAM 1 KO animals (Figure 7A; compare with Figure 2C). We collected MUA and VEP responses of non-deprived PV-Cre animals injected at P14 with AAV delivering the shRNA against SynCAM 1 and scrambled sequence (Figure 7B; Knockdown/KD (PV-Cre^AAV-shCadm15^) and Control/Ctrl (PV-Cre^AAV-shScramble^). No detrimental effects of the viral injection were observed, and these AAV-shScramble, Control (Ctrl) injected animals had a C/I ratio and VEP amplitudes almost identical to non-deprived and non-injected animals during the critical period (Figure 7C, compare with Figure 2D; Ctrl ND C/I Ctrl=2.3±0.2; Contra/right eye=163±19.7 μV, Ipsi/left eye=72±10.2 μV; Supplementary Table 3). No gross changes in visual responses were observed when knockdown of SynCAM 1 was targeted to PV^+^ interneurons (Figure 7C and 7D, KD ND C/I=2.1±0.2; Contra/right eye=162±10.2 μV, Ipsi/left eye=78±4.6 μV; Supplementary Table 3). Short monocular deprivation from P60-P64 had no effect on the C/I ratio of adult control mice, as expected (Figure 7C and 7D; Ctrl MD C/I=2±0.2; Contra/closed eye=171±6.6 μV, Ipsi/open eye=87±7.1 μV; Supplementary Table 3). Importantly, knockdown of SynCAM 1 in PV^+^ interneurons resulted in robust plasticity in adult mice after short, 4-day monocular deprivation (Figure 7C and 7D; KD MD C/I=1.3±0.03, p=0.002; Supplementary Table 3). This plasticity was due to a significant depression of closed eye responses, similar to plasticity of WT mice during the critical period (Figure 7D, compare with Figure 2E; KD MD Contra/closed eye=114±10.5 μV, p=0.039; Ipsi/open eye=90±8.9 μV, p=ns; Supplementary Table 3).

**Figure 7.**
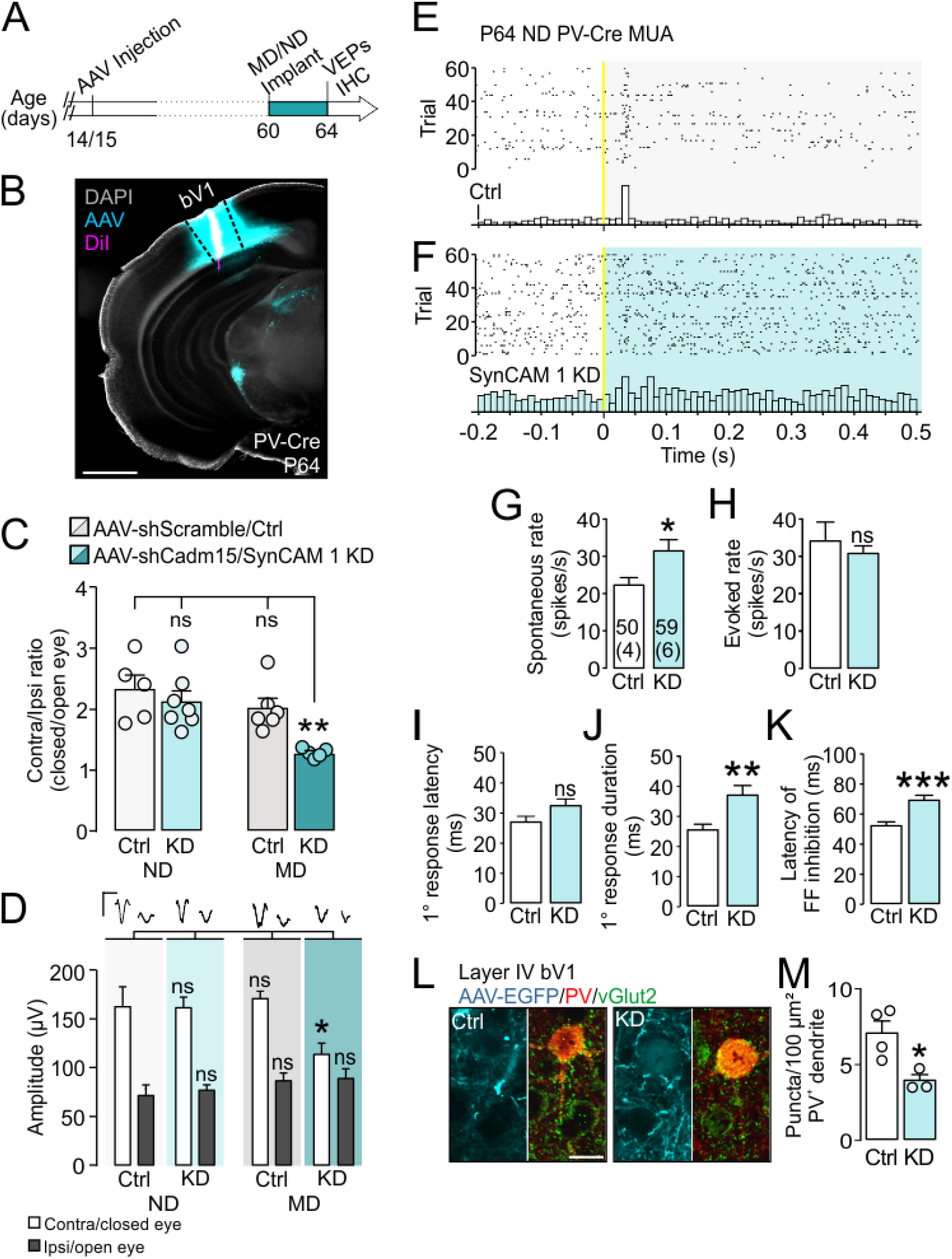
PV^+^ interneuron-specific knockdown of SynCAM 1 in V1 extends the critical period. A)Timeline of the experimental approach. AAV-shScramble or AAV-shCadm15 delivering shRNA for SynCAM 1 knockdown (KD) were injected into left bV1 of PV-Cre mice at P14/15. For recordings, titanium headposts were implanted at P60 and right eyelids were sutured shut in randomly selected animals. Non-sutured mice served as controls. 4 days later, the right eye was reopened and VEPs were recorded. For immunohistochemistry, non-deprived animals were sacrificed at P64 without prior headpost implantation.
B)Falsely colored representative section of an AAV-injected animal depicting localized viral spread and a representative electrode tract through the injection site. Some viral transport was apparent in the subcortical regions, but no AAV-positive fibers were seen entering the LGN. Scale bar: 1 mm
C)C/I ratios of non-deprived shScramble-injected control animals and animals with shCadm15-mediated SynCAM 1 knockdown in PV^+^ interneurons (KD) were almost identical at P64 (also compare with Figure 2D). 4 days of MD at P60 did not affect shScramble-injected control animals, as expected. In contrast, MD from P60-64 robustly decreased the C/I ratio in KD mice, indicating that plasticity occurs in adult mice upon targeting SynCAM 1 in PV^+^ cells.
D)Visual responses of naïve, non-deprived AAV-shScramble (Ctrl) and AAV-shCadm15 (KD) injected animals were almost identical and similar to naïve P28 animals (compare with Figure 2E). MD had no effect on shScramble injected control animals as expected, but it significantly depressed closed eye responses after SynCAM 1 knockdown in PV^+^ cells. Representative VEPs collected from Ctrl and KD mice are shown on top. Scale bars: 200 μV and 0.2 s. Panels C, D: ns, not significant; *, p<0.05; **, p<0.01; One-way ANOVA. Data presented as mean±SEM See Supplementary Table 3 for details.
E,F) Representative raster plots and PSTHs of MUA recorded from layer IV of P64 control (E) or SynCAM 1 KD PV-Cre mice (F). Stimulus onset is indicated in yellow. Scale bars: 5 spikes/s. G-H) Spontaneous firing rate was significantly increased after SynCAM 1 KD in PV^+^ interneurons (G), while the poststimulus evoked rate was not significantly different between the groups (H). I-K) Latency of primary response was not significantly increased after SynCAM 1 KD (I), but the duration was significantly increased (J). Feed-forward inhibition was significantly delayed after SynCAM 1 KD (K). Panels G-K: ns, not significant; *, p<0.05; **, p<0.01; ***, p<0.001; Mann-Whitney rank sum test. Data presented as mean±SEM. See Supplementary Table 4 for details.
L) Immunofluorescence labeling of vGlut2^+^ terminals in bV1 in AAV-shScramble (Ctrl) and AAV-shCadm15 (KD) injected PV-Cre animals. Single optical confocal sections containing dendritic segments near the injection site and devoid of EGFP signal were analyzed by immunostaining for PV (red) and vGlut2 (green). GFP (cyan) marks Cre^-^/non-PV^+^ cells, in which no shRNA expression occurred. Scale bar: 15 μm. M) Quantification of data as in (M) shows that KD of SynCAM 1 significantly reduced the density of vGlut2^+^ TC terminals onto PV^+^ dendrites in V1. vGlut2 puncta density in control, shScramble injected animals was lower than in non-injected WT animals (Figure 4) due to different antibody titers of lots used. *, p<0.05;unpaired t-test. Data presented as mean±SEM, Ns are indicated.

We observed an increase in MUA firing rate in layer IV, reflecting the expected developmental increase in neuronal firing rates in adult (P64) animals compared to critical period mice (Figure 7E-G, compare with Figure 5B-D; Supplementary Table 4) (Chen et al., 2015). However, the spontaneous firing rate was even higher after SynCAM 1 knockdown in PV^+^ interneurons (Figure 7F and 7G; Ctrl=23±1.7 Hz, KD=32±2.8 Hz, p=0.032; Supplementary Table 4), indicating disinhibition (Atallah et al., 2012; Gu et al., 2013; Kaplan et al., 2016). The magnitude and latency of visual response was not significantly affected after SynCAM 1 knockdown (Figure 7H and 7I; evoked firing rate Ctrl=34±4.8 Hz, KD=31±2 Hz; primary response latency Ctrl=27±1.6 ms, KD=33±2.1 ms; Supplementary Table 4). However, SynCAM 1 KD mice had significantly protracted responses and delayed feed-forward inhibition, as observed in SynCAM 1 KO mice (Figure 7J and 7K, compare with Figure 5G and 5H; primary response duration Ctrl=26±1.6 ms, KD=38±3 ms; p=0.007; latency of feedforward inhibition Ctrl=53±2.1 ms, KD=71±3 ms, p<0.001; Supplementary Table 4). Knockdown of SynCAM 1 in cortical PV^+^ interneurons is hence sufficient to maintain V1 in an immature state and extend plasticity beyond the critical period.

Does disinhibition of V1 in the cell-type and region-specific knockdown of SynCAM 1 in PV^+^ interneurons share a common cellular mechanism with the global loss of SynCAM 1? To address this question, we quantified the recruitment of TC terminals onto PV^+^ dendrites in V1 of adult control and SynCAM 1 KD mice near the injection sites of Control and KD constructs (Figure 7L and 7M). PV-specific knockdown of SynCAM 1 in V1 reduced the density of vGlut2^+^ terminals in contact with PV^+^ dendrites by 45% (Figure 7L and 7M; Ctrl=7±0.5 puncta/100 μm^2^, N=4 animals; KD=4±0.3 puncta/100 μm^2^, N=3 animals; p=0.026, t=3.1, df=5). Intracortical inputs to PV^+^ interneurons did not change, as the density of vGlut1 puncta in contact with PV^+^ dendrites remained intact after SynCAM 1 knockdown (Ctrl=34±1.7 puncta/100 μm^2^; KD=31±7.7 puncta/100 μm^2^; data not shown). Puncta size was unaltered for both vGlut1 and vGlut2 across conditions (data not shown). The synaptic maturation of the thalamocortical visual circuit hence engages cell-autonomous, postsynaptic, and input-specific function of SynCAM 1 as synapse organizer in cortical PV^+^ interneurons. These results demonstrate the central role of SynCAM 1-dependent feed-forward TC afferents onto PV^+^ interneurons to control the closure of the critical period for visual deprivation.

## Discussion

Our findings suggest that the closure of the visual critical period requires SynCAM 1 in cortical PV^+^ interneurons to organize the thalamocortical synaptic inputs. Five lines of evidence confirm SynCAM 1 as a novel PV^+^-autonomous factor that controls the maturation of inhibitory drive and developmental plasticity in the cortex. First, visual activity selectively regulates SynCAM 1 protein levels in the visual cortex. Second, SynCAM 1 acts in PV^+^ interneurons in V1 to selectively control excitatory feed-forward TC inputs, but not local intracortical synaptic connectivity. Third, PV^+^ interneurons are structurally immature in the absence of SynCAM 1 as apparent in reduced PNN deposition. Fourth, critical period closure requires the cell-autonomous, SynCAM 1- mediated maturation of feed-forward, PV^+^-driven inhibition. Fifth, short visual deprivation elicits robust plasticity in SynCAM 1-deficient circuits, including adult animals. These conclusions are supported by results from KO mice lacking SynCAM 1 and from mice in which this protein was selectively knocked down in PV^+^ interneurons. We propose that SynCAM 1 functions as a synaptic brake on cortical plasticity by controlling the number of thalamocortical inputs onto PV^+^ interneurons.

SynCAMs, neuroligins, and other synaptic adhesion molecules modulate and can even instruct synapse formation and plasticity throughout the central nervous system (Missler et al., 2012; Schreiner et al., 2017; Shen and Scheiffele, 2010; Frei and Stoeckli, 2017). Their roles in excitatory synapse formation and maturation in pyramidal neurons are well described, but their functions in other cell types are only beginning to be defined (Chen et al., 2017; Park et al., 2016; Polepalli et al., 2017; Rathjen et al., 2017). Neuroligin-3 controls both metabotropic and ionotropic glutamate receptor signaling in hippocampal CA1 PV^+^ interneurons (Polepalli et al., 2017). Unlike neuroligin-3, which controls the functional maturation of PV^+^ interneuron inputs, we found that PV^+^-specific loss of SynCAM 1 reduces excitatory synapse number to PV^+^ interneurons. Moreover, SynCAM 1 selectively controls TC inputs, but not intracortical inputs, identifying synapse-type specific roles of this molecule. Expression of SynCAM 1 is not restricted to thalamorecipient layers, indicating that a presynaptic partner may confer the synapse type-specific roles of SynCAM 1 we report here. Both homophilic and heterophilic interactions between SynCAM 1 and other adhesion molecules across the synaptic cleft are thought to underlie its synapse-organizing roles (Fogel et al., 2007; Perez de Arce et al., 2015). The trans-synaptic molecular partners for SynCAM 1 in the TC circuit are as yet unknown, but are worthy of future studies.

Other mechanisms that control excitatory drive onto PV^+^ interneurons include Neuregulin-1/ErbB4 signaling (Fazzari et al., 2010; Sun et al., 2016), pentraxins (Chang et al., 2010; Gu et al., 2013; Pelkey et al., 2016) and the NogoR receptor (Stephany et al., 2016). Interestingly, all three are critical for ocular dominance plasticity (ODP), suggesting a pivotal role of excitatory drive onto PV^+^ interneurons in the initiation of ODP (Gu et al., 2013; Kuhlman et al., 2013; McGee et al., 2005; Sun et al., 2016). Specific synapse types that are affected by these proteins remain unclear. It is currently also unknown if these pathways cross-talk with SynCAM 1 signaling in V1. A recent study reported on an interplay between SynCAM 1 and Neuregulin-1/ErbB4 signaling in hippocampal PV^+^ interneurons (Yamada et al., 2013). However, Neuregulin-1 expression levels are similar between WT and SynCAM 1 KO mice (data not shown). Further, a physical association between SynCAM 1 and Narp or NogoR appears unlikely as SynCAM 1 is only known to preferentially interact with other trans-synaptic adhesion molecules (Fogel et al., 2007; Perez de Arce et al., 2015). SynCAM 1 is often compared to another postsynaptic organizer, neuroligin-1 (Czondor et al., 2013; Sara et al., 2005). Similar to neuroligin-1, SynCAM 1 protein is abundant in all cortical layers (Figure 1) (Singh et al., 2016). However, the loss of neuroligin-1 and its astrocyte-secreted, synaptic bridge protein Hevin affects both local, vGlut1^+^ and TC, vGlut2^+^ inputs (Risher et al., 2014; Singh et al., 2016), while here we demonstrated that the loss of SynCAM 1 selectively affects TC afferents. Moreover, SynCAM 1 acts in PV^+^ interneurons to control their TC inputs, making it the first synapse-organizing molecule known to have this property.

Well-known molecular brakes that broadly control developmental plasticity, such as PNNs and myelin (Lensjo et al., 2017; Pizzorusso et al., 2002; McGee et al., 2005), ensure the stabilization of synapses after extensive remodeling during the critical period. They control PV^+^ interneuron function on multiple levels that converge on interactions of PV^+^ interneurons with the extracellular matrix (ECM) and PNNs. Specifically, transcriptional programming initiated by non-autonomous transcription factor Otx2 opens the critical period in V1 and leads to tight structural integration of PV^+^ interneurons into the surrounding ECM (Sugiyama et al., 2008; Sugiyama et al., 2009). PirB and myelin-receptor NogoR are molecular brakes that interact with each other, and NogoR binds to components of the ECM (Atwal et al., 2008; McGee et al., 2005). Age-dependent deposition of PNNs is pivotal for closure of the critical period (Hou et al., 2017; Miyata et al., 2012) and removal of PNNs from adult cortex reinstates immature network properties, including heightened plasticity, in V1 (Lensjo et al., 2017; Miyata et al., 2012; Pizzorusso et al., 2002). We found that PNN density in SynCAM 1 KO mice is lowered at all developmental stages and even in the adult did not reach WT critical period levels. Early steps of PV^+^ interneuron maturation likely proceed normally in SynCAM 1 KO mice, since PNNs are still present, though they are structurally compromised. This is consistent with normal critical period opening and the recruitment of Otx2 in these mice. While a physical association between SynCAM 1 and PNN components has not been described, the reduced PNN formation in the KO mice may be due to decreased feed-forward excitation onto SynCAM 1-deficient PV^+^ interneurons which could negatively impact the activity-dependent formation of PNNs (Dityatev et al., 2007).

Although less numerous than intracortical synapses onto PV^+^ cells, TC synapses are much stronger so even a small reduction in their density can result in circuit disinhibition (Cruikshank et al., 2007; Ji et al., 2016; Kloc and Maffei, 2014; Miyata et al., 2012). Consequently, the SynCAM 1-dependent reduction of TC inputs results in profound hyperexcitation, disinhibition and a protracted critical period, in agreement with the role of cortical inhibition in shaping the ODP (Figure 7) (Kuhlman et al., 2013; Trachtenberg, 2015). Previous studies demonstrated increased synaptic plasticity in the absence of SynCAM 1 in multiple brain regions (Rathjen et al., 2017; Robbins et al., 2010). Our work in the cortex, this work now reveals a synapse- and circuit specific mechanism for SynCAM 1-mediated restriction of network plasticity. The normal gross visual responses, but absent critical period closure in both global SynCAM 1 KO and in PV-Cre SynCAM 1 KD mice highlight the selective contributions of trans-synaptic interactions to circuit maturation. We also noted interesting differences in visual responses of these two models. The differential plasticity of closed (contra) and open (ipsi) eye pathways in SynCAM 1 KO and SynCAM 1 KD mice that lead to the same effect on ODP likely reflects additional roles of SynCAM 1 in other cortical cell and synapse types, reminiscent of the multiple roles of NogoR (McGee et al., 2005; Stephany et al., 2014; Stephany et al., 2016). In young animals, excitatory synapses on pyramidal neurons are critical for open eye potentiation (Ranson et al., 2012) and specific contributions of SynCAM 1 to the plasticity of closed *vs* open eye pathways remain to be studied. SynCAM 1 KO mice have a reduced frequency of excitatory miniature events in the hippocampus (Robbins et al., 2010) and it is possible that such reduced baseline excitability dampens the effects of cortical disinhibition and accounts for the differences in spontaneous and evoked activity between the SynCAM 1 KO and SynCAM 1 KD mice. Retinal input is the main driver of primary visual response (Shen and Colonnese, 2016), and the late MUA response in SynCAM 1 KO animals likely reflects the previously reported, modest delay in retinal visual processing in these mice (Ribic et al., 2014). Consistent with this interpretation, MUA responses have normal latency in SynCAM 1 KD animals. More generally, phenotypic differences between global and conditional KO mouse models are not uncommon (Chen et al., 2017; Polepalli et al., 2017; Stephany et al., 2014; Sommeijer et al., 2017) and in our case are likely due to a role for SynCAM 1 in cells other than PV^+^ interneurons. Importantly, the disinhibition of V1 evident in prolonged spiking activity and retarded feed-forward inhibition, is almost identical between the two models, highlighting the specific requirement for SynCAM 1 at the postsynaptic side of TC inputs to PV^+^ cells. Both our mouse models of diminished SynCAM 1 expression therefore support a requirement for reduced inhibitory drive to permit plasticity in the adult visual cortex (Harauzov et al., 2010; Kuhlman et al., 2013). The increased expression of SynCAM 1 in adult V1 matches its role as a synaptic brake that restricts adult plasticity by maintaining strong TC inputs onto PV^+^ cells. The increase of SynCAM 1 in the deprived hemisphere of the developing V1 observed after MD may reflect a response to maintain homeostasis and limit remodeling. ODP still proceeds despite SynCAM 1 upregulation in WT mice, but higher SynCAM 1 may contribute to keeping the response of the actively remodeling circuits within a physiologically relevant limit.

Our results demonstrate that SynCAM 1 is novel cell-autonomous factor that controls cortical plasticity and the development of feed-forward, excitatory thalamic inputs to V1. SynCAM 1 acts as a synaptic switch for the maturation of cortical inhibition, supporting the wiring and proper balance of the TC circuit for vision. Our results, as well as unbiased transcriptome and proteome screens (Loh et al., 2016; Lyckman et al., 2008), demonstrate that SynCAMs are the only trans-synaptic organizers known to be both exclusive to excitatory synapses and regulated by visual activity. These results therefore provide support for specific and non-redundant roles of synapse-organizing molecules in circuit development *in vivo*. Our study also provides evidence that the maturation of TC-driven, feed-forward cortical inhibition is a necessary step for the closure of the critical period. Recent findings suggest that inhibitory networks in the visual thalamus can also modulate ocular dominance plasticity (ODP) in the cortex (Sommeijer et al., 2017). Thalamic axons themselves display plasticity after monocular deprivation, raising the question of specific contributions of cortical PV^+^ interneurons to critical period plasticity (Jaepel et al., 2017; Sommeijer et al., 2017). Our study demonstrates that excitatory, thalamocortical (TC) inputs to PV^+^ interneurons in V1 are essential for critical period closure, in agreement with a central role of cortical inhibition in critical period regulation (Gu et al., 2013; Kuhlman et al., 2013; Stephany et al., 2016; Sun et al., 2016). The identification of SynCAM 1 as a novel, PV^+^-autonomous molecular brake on plasticity highlights that distinct mechanisms control critical period opening *vs* critical period closure, and sheds light on the profound impacts of excitatory/inhibitory imbalance and regulatory feedback loops that are frequently implicated in neurodevelopmental disorders (Ebert and Greenberg, 2013; Mullins et al., 2016; Nelson and Valakh, 2015; Zoghbi and Bear, 2012).

## Author contributions

Conceptualization, A.R.; Methodology: A.R. Software: A.R. Investigation: A.R.; Writing – Original Draft, A.R.; Writing–Review & Editing, A.R. and T.B.; Funding Acquisition, A.R., M.C.C. and T.B.; Resources, M.C.C. and T.B.; Supervision, T.B.

## Acknowledgments

We thank the members of the Biederer and Crair laboratories for valuable feedback and assistance, esp. Drs. O.S. Dhande, T.J. Burbridge and E.J. Mohns for discussions and experimental guidance in the early stages on of the project and Bea Carbone for preparation of neuronal cultures. We are grateful to Dr. T. Momoi for generously providing the SynCAM 1/RA175 KO mouse line, Drs. M. Picciotto and E. Wohleb for providing AAV vectors, Dr. L. Reijmers for providing the PV-Cre mouse line, the CED support team for programming advice and all unnamed colleagues for constructive discussions. This work was supported by the National Institute of Health Grants R01 DA018928 (to T.B.), R01EY015788 (to M.C.C.), R01EY023105 (to M.C.C.), U01NS094358 (to M.C.C.), P30EY026878 (to M.C.C.), and the Knights Templar Eye Foundation (to A.R).

## STAR Methods

Further information and requests for resources and reagents should be directed to and will be fulfilled by the Lead Contact, Thomas Biederer.

### Experimental Model and Subject Details

#### Animals

Experiments were performed on C57BL6/J wild type mice (The Jackson Laboratory, Bar Harbor, ME), SynCAM 1 KO mice (Fujita et al., 2006) and their wild type littermates, and heterozygous PV-Cre mice (JAX 008069; (Hippenmeyer et al., 2005; Kuhlman et al., 2013)). All mouse lines were maintained on a C57BL6/J background and KO mice had been backcrossed more than 10 times. Animals of both sexes from postnatal day 7 (P7) to P70 were used for all experiments as indicated below and stated in the figure legends. Littermates were compared in all experiments and the experimenter was blind to the genotype of animals used. Animals were kept on a 12/12 hour light/dark cycle with food and water *ad libitum*. All experiments were performed during the light phase (7 AM-7 PM). For neuronal cultures, pregnant Sprague-Dawley rat dams were purchased from Charles River Laboratories (Wilmington, MA). Animals were treated in accordance with Institutional Animal Care and Use Committee guidelines.

### Method Details

#### Antibodies

Primary antibodies and their properties are listed in Key Resources Table. For all immunostainings, secondary antibodies were applied in the absence of primary antibodies as a control. Secondary antibodies and reagents are listed in Key Resources Table.

#### Tissue preparation for biochemistry and microscopy

Animals were anesthetized with ketamine (100 mg/kg) and xylazine (10 mg/kg) in saline. For protein isolation (animals aged P7-P45 for Fig. 1A, P28 for Fig. 1C and 1D), visual cortices was isolated according to stereotactic coordinates (0.5-1 mm anterior to λ, 2-3 mm lateral to midline) followed by sonication in 8 M urea. For GABA and glutamate receptor immunoblots, crude synaptoneurosomes were prepared as described (Villasana et al., 2006). Protein concentrations were determined using the BCA method (ThermoFisher Scientific, Holtsville, NY). For microscopy, animals (P21-P70, as indicated in figure legends) were transcardially perfused first with ice cold PBS and then with 4% PFA (in PBS, pH 7.4). Brains were isolated and postfixed overnight in 4% PFA and washed overnight in PBS (all at 4°C). Brains were then embedded in 3% agarose in PBS and sectioned at 40-60 μm using vibrating microtome (Leica VT1000, Leica Biosystems, Nussloch, Germany or Vibratome 1500, Harvard Apparatus, Holliston, MA). Sections were stored in PBS with 0.01% sodium azide (Sigma Aldrich, St. Louis, MO) at 4°C.

#### Quantitative immunoblotting

Proteins from cortical homogenates or crude synaptosomes (10–30 μg for V1 and 60 μg for LGN, prepared as described above) were subjected to immunoblotting using standard procedures (Fogel et al., 2007) and scanned with either Odyssey Infrared Imaging System (LI-COR Biosciences, Lincoln, NE) or FluorChem M (Protein Simple, San Jose, CA). Antibodies used are listed in Key Resource Table. YUC8 and 3E1 provided almost identical signal in blots, except that SynCAM 1 signal in the LGN was better visible with YUC8, likely due to its higher affinity for different glycosylation states of SynCAM 1 (Fogel et al., 2007). For all blots imaged using FluorChem M, milk was replaced with BSA (Sigma) for blocking and probing. For quantitative immunoblotting in Figure 1, secondary IRDye800 antibodies or anti-IgG Alexa 647 were used. Quantification was performed using the gel analysis plugin in ImageJ, where actin served as loading control for all samples.

#### Culturing and immunolabeling of primary neurons

Cortical neurons were prepared from rats at E18 as described (Biederer and Scheiffele, 2007) with modifications. In brief, dissected cortices were incubated in 0.05% trypsin at 37 °C for 20 minutes (Invitrogen, Carlsbad, CA; 25300054) and plated at a density of ~30,000 cells per coverslip. Dissociated cells were plated on poly-l-lysine (Sigma P1274) and incubated in a cell culture incubator with 5.0% CO2. Cytosine arabinoside (Sigma C1768) was added at a final concentration of 2 μM per well 2 days in vitro to prevent glia cell overgrowth. Cells were washed with ice-cold PBS and fixed at DIV 7 and DIV 14 in ice-cold 4% PFA/4% sucrose for 15 minutes, permeabilized with 0.1% Triton-X100 in PBS for 10 minutes at RT and blocked in 5% FBS in PBS for 1 hour at RT. Coverslips were later sequentially incubated for 1 hour at RT in anti-SynCAM 1, anti-Parvalbumin and WFA (see Key Resource Table for more details) and their corresponding secondary antibodies. All antibodies were diluted in PBS and coverslips were washed 3 × 10 minutes in PBS at RT in between all antibody incubations. Coverslips were mounted with Aqua-Mount (Polysciences Inc., Warrington, PA) and imaged as described below.

#### Immunohistochemistry and confocal microscopy

Primary antibodies used in double- and triple-labeling experiments were applied sequentially and blocking steps were performed using normal horse serum. Visual cortex sections were first washed in PBS and non-specific antibody binding sites were blocked with 3% normal serum and 0.03% Triton-X 100 (Sigma) in PBS for 1 h at RT. Primary and secondary antibodies were diluted in 3% normal serum and 0.03% Triton-X 100 in PBS and incubated either for 24–48 hours at 4°C (primary antibodies) or 1 hour at room temperature (secondary antibodies). After the antibody incubation steps, sections were washed in PBS and floated on slides in distilled water before coverslipping with mounting medium (Aqua-Polymount, Polysciences Inc., Warrington, PA, USA). Confocal microscopy was performed on a Leica TCS SPE DM2500 microscope or a Leica TCS SP8. Images were acquired with ACS AP 40x oil lens with 1.15 NA for Figures 1, 3 and Supplementary Figure 3 or ACS APO 63x oil lens with 1.3 NA for Figures 4, 7 and Supplementary Figures 2 and 5 using identical settings for each group within an experiment. All images were acquired in binocular V1, layer IV. Low magnification images were acquired with Zeiss Axio Scope (Carl Zeiss, Jena, Germany) or BZ-X700 (Keyence, Osaka, Japan).

*Image quantification*

For Figures 3 and Supplementary Figure 3, Otx+, WFA^+^ and PV^+^ cells were counted manually using ImageJ (NIH, Bethesda, MD). For quantification, only single optical sections were used, except for Supplementary Figure 3 where maximum intensity projection images were used and for analysis of WFA particle density, where Pipsqueak plugin used sum intensity projection. Quantification of WFA^+^ puncta density was performed using Pipsqueak plugin for ImageJ (Slaker et al., 2016). Quantification of SynCAM 1 and NeuN in cortex was performed with ImageJ measuring the % area covered by signal. SynCAM 1 signal was normalized to NeuN values to normalize for slight variations in section thickness due to vibratome sectioning. Quantification of vGlut1, vGlut2 and SynCAM 1 puncta was performed as previously described (Park et al., 2016). Briefly, contours of PV^+^ dendrites in layer IV were manually outlined. vGlut1, vGlut2 and SynCAM 1 images were thresholded, binarized and the density of puncta in contact with PV^+^ dendrites (ROI) was counted using particle analyzer tool with a vGlut1 and SynCAM 1 cutoff of0.1 μm^2^ and vGlut2 cutoff of 0.2 μm^2^. On average, 10–20 dendritic segments were collected from each animal from 3–6 brain sections. For SynCAM 1 knockdown validation *in vivo*, ROI was defined as PV^+^ cell body and 20 cells on average were analyzed per animal for both SynCAM 1 KD/PV-Cre^AAV-shCadm15^ and Control/PV-Cre^AAV-shScramble^. All values were averaged per animal before final statistical analysis.

#### Bulk anterograde labeling and quantification of eye-specific segregation

Retinal ganglion cell projections from the right and the left eye were bulk labeled with CTB Alexa-488 and CTB-Alexa 555. The tracer was diluted to 1 mg/ml in 0.9% saline. At P12/13, mice were anesthetized and injected with 1-2 μl tracer per eye using a glass pulled pipette and Nanoject (Drummond Scientific, Broomall, PA). 48 hours later mice were transcardially perfused and the brains were fixed overnight in 4% PFA. Coronal sections (80 μm thickness) were collected with a vibratome as described above, mounted in Aquamount and imaged with a CCD camera (Zeiss). Analysis of segregation of contralateral and ipsilateral projections in dLGN was performed as previously described (Torborg and Feller, 2004). Briefly, images were background subtracted with a rolling ball radius of 200 in ImageJ, and the three sections with the largest ipsilateral (Alexa 555 labeled) area were used for analysis. The logarithm of the intensity ratio, R=log10 (ipsilateral channel fluorescence intensity/contralateral channel fluorescence intensity), was determined for each pixel, and a segregation index for each animal was computed as the mean of the variance of the distribution of R values. A larger segregation index (higher variance) is indicative of better segregation (Torborg and Feller, 2004).

#### AAV cloning, packaging, purification and shRNA validation

For SynCAM 1 knockdown *in vivo*, sequence shCadm15 (Faraji et al., 2012) was cloned into pAAV-dsRed-Sico-shRNA (Wohleb et al., 2016) (kindly provided by Dr. Marina Picciotto, Yale University). 70% confluent AAV-HEK293 cells (Agilent, Santa Clara, CA) were transfected with pHelper, AAV/DJ Rep-Cap and pAAV-dsRed-Sico-shCadm15 or pAAV-dsRed-Sico-shScramble using PEI method (Sonawane et al., 2003). Cells were collected after 72 hours and AAV was purified using iodixanol gradient (Hermens et al., 1999). AAV was further concentrated using Amicon 15 (EMD Milipore Sigma). Titer was determined as in (McClure et al., 2011). 600 nl of virus (3 × 10^12^ GC/ml) was injected at 1 nl/s into layer 4 of bV1 (~350 μm depth) at P14 using stereotaxic apparatus (Stoelting, Wood Dale, IL) and glass pipette attached to Hamilton syringe (Hamilton Robotics, Reno NV) using micro-syringe pump (Micro4, WPI, Sarasota, FL) at following coordinates: 0-1 mm anterior to λ, 2.5–3 mm lateral to midline (Valverde, 1998). For targeting validation *in vitro*, shCadm15 sequence was cloned into pSico (Ventura et al., 2004). Confluent HEK293 cells were transfected with pCAGGS-SynCAM 1 (Stagi et al., 2010), pSico-Cadm15 and pAAV-GFP-Cre (kindly provided by Dr. Dong Kong, Tufts University) using PEI transfection. Cells were collected 72 hours later and lysed in RIPA buffer. 30 μL of protein homogenate was immunoblotted for SynCAM 1 and quantified as described above. For targeting validation in cultured neurons, primary cortical cultures were transfected at DIV 5 as above using Lipofectamine (Invitrogen) and fixed at DIV 14. SynCAM 1 signal was quantified as described above. For targeting validation *in vivo*, animals injected with pAAV-dsRed-Sico-shCadm15 or pAAV-dsRed-Sico-shScramble were perfused as described above. SynCAM 1 immunohistochemistry was described as above and puncta were quantified using PV^+^ signal as ROI (as described above).

#### Eyelid suture

Mice were anesthetized with isoflurane in oxygen (2% induction, 1.0-1.8% maintenance) and placed under a surgical microscope. Area around the right eye was sterilized with alcohol swabs and lid margins were trimmed. Three mattress stitches were placed using 7–0 nylon sutures and the lids were further attached using VetBond (3M). After that, ophthalmic antibiotic ointment was applied to the suture. Mice were monitored daily for the integrity of the sutures and signs of infection. Animals whose eyelids were not fully sutured and animals that removed their sutures were excluded from further experiments. At the end of the deprivation period, after the head-plate implantation (see below), the stitches were removed, and lid margins separated. Eyes were flushed with sterile saline and checked for clarity. Mice with corneal opacities, cataracts or signs of infection were excluded from further study.

#### In vivo electrophysiology

Recordings were performed on awake female and male mice, ages P21 to P64, using spherical treadmill as described in (Niell and Stryker, 2010). 4–7 days before the recording session, custom made titanium or aluminum (for precritical period mice) head-plate implants were cemented to the mouse skull. Animals were anesthetized with isoflurane in oxygen (2% induction, 1.0-1.8% maintenance), warmed with a heating pad at 38°C and given subcutaneous injections of Buprenorphine SR (1mg/kg) and 0.25% Bupivacaine (locally). Eyes were covered with Puralube (Decra, Northwich, UK). Scalp and fascia from Bregma to behind lambda were removed, and the skull was cleaned, dried and covered with a thin layer of cyanoacrylate (VetBond; 3M, Maplewood, MN) before attaching the head plate with dental cement (RelyX, 3M). The well of the head plate was filled with silicone elastomer (Reynold Advanced Materials, Brighton, MA) to protect the skull before recordings. Animals were single housed after the implantation and monitored daily for signs of shock or infection. 1-2 days before the recording, the animals underwent 1–2 20–30 minutes handling sessions and 1-2 10-20 minutes session in which the animals were habituated to the spherical treadmill (Dombeck et al., 2007). On the day of recording, the animals were anesthetized as above and small craniotomies (~0.5 mm in diameter) with 18G needles were made above bV1 and cerebellum. The brain surface was covered in 2–3% low melting point agarose (Promega, Madison, WI) in sterile saline and then capped with silicone elastomer. Animals were allowed to recover for 2–4 h. For the recording sessions, mice were placed in the head-plate holder above the free-floating ball and allowed to habituate for 5–10 minutes. The agarose and silicone plug were removed, the reference insulated silver wire electrode (A-M Systems, Carlsborg, WA) was placed in cerebellum and the well was covered with warm sterile saline. A multisite electrode spanning all cortical layers (A1 × 16-5mm-50–177-A16; Neuronexus Technologies, Ann Arbor, MI) was coated with DiI (Invitrogen) to allow *post hoc* insertion site verification and then inserted in the brain through the craniotomy. The electrode was lowered until the uppermost recording site had entered the brain and allowed to settle for 20–30 minutes, after which the ipsilateral eye response was checked to confirm the proper location in V1. The well with the electrode was then filled with 3% agarose to stabilize the electrode and the whole region was kept moist with surgical gelfoam soaked in sterile saline (Pfizer, MA). Minimum 2 penetrations were made per animal to ensure proper sampling of the craniotomy. Recordings sessions typically lasted 2–3 h. After the recording, mice were euthanized with an overdose of ketamine and xylazine. The brains were then isolated and fixed in 4% PFA overnight at 4°C. Brains were subsequently sectioned at 100 μm using a vibrating microtome. The sections were incubated in DAPI (Sigma), floated on slides and mounted in Aquamount. The sections were imaged on a Keyence microscope as described above to confirm the electrode location within bV1.

#### Visual stimuli, data collection and analysis

Visual stimuli were generated with MATLAB (MathWorks, Natick, MA) using the Psychtoolbox extension (Brainard, 1997) and Spike2 (CED, Cambridge, UK). Varying frequencies and orientations of M-field sinusoidal gratings at 100% contrast were displayed on a gamma corrected 17” LCD (Niell and Stryker, 2008) for 1.5 s with 0.2 s interstimulus interval (grey screen). Stimuli were presented in randomized fashion and each stimulus was presented 30–50 times on average during a recording session. The screen was centered 20 cm from the mouse’s eye, covering ~80° of visual space. 30–50 light-emitting diode (LED) flashes were presented before or after the sinusoidal gratings with 10 s interstimulus interval. Visual responses signals were preamplified 10x (MPA8I preamplifiers; Multi Channel Systems MCS GmbH, Reutlingen, Germany) and then fed into a 16-channel amplifier (Model 3500; A-M Systems), amplified 200x and band-pass filtered 0.3-5000 Hz. The signals were sampled at 25 kHz using Spike2 and data acquisition unit (Power 1401-3, CED). Stationary and movement stages of animal behavior were separated using an optical mouse that tracked the movement of the styrofoam ball and was interfaced with LabView (Austin, TX) and Spike2, or using video recordings timed to stimuli presentation (WansView, Shenzhen, PRC). Only stationary, non-running stages were analyzed offline using Spike2 software (CED). LFPs were analyzed as waveform averages, triggered by stimulus onset. Visually evoked potentials (VEPs) were defined as negative-going events occurring within 200 ms following stimulus onset, having an amplitude of more than 3x standard deviation and having a width at half maximum of less than 50 ms (Li et al., 2013). For sinusoidal gratings, frequency evoking a maximum response in test animals was analyzed (0.15 Cycles per degree at all orientations). Signals evoked by sinusoidal gratings and LEDs were identical in nature, so their amplitude values were averaged. For multiunit analysis, spikes were extracted from band-pass filtered data using thresholds (3x standard deviation) and sorted in Spike 2. Peri-stimulus time histogram (PSTH) analysis was performed with Spike2 using 0.01 s bins. Spontaneous firing was calculated as average firing rate before stimulus presentation with 0.2 s offset. Spontaneous firing was subtracted from peak poststimulus firing rate to determine evoked firing rate. Spectral analysis was performed on raw LFP traces as previously described (Mohns and Blumberg, 2008), with gamma-band oscillations defined as 40-70 Hz, as described in Chen et al., 2015.

### Quantification and Statistical Analysis

All quantitated analyses were performed with the researcher blind to the condition. Statistical analyses were performed in SigmaPlot 11 and 13 (San Jose, CA) or GraphPad Prism 7.0 (GraphPad Inc., La Jolla, USA) using t-test and one or two-way ANOVA with post-hoc comparisons (as indicated in text, figure legends and Supplementary Tables), unless stated otherwise. When comparing two independent groups, normally distributed data were analyzed using a Student’s t-test. In the case data were not normally distributed a Mann-Whitney rank sum test was used. All data are reported as mean ± SEM, where N represents number of animals used, unless indicated otherwise. Target power for all sample sizes was 0.8. In all cases, alpha was set to 0.05.

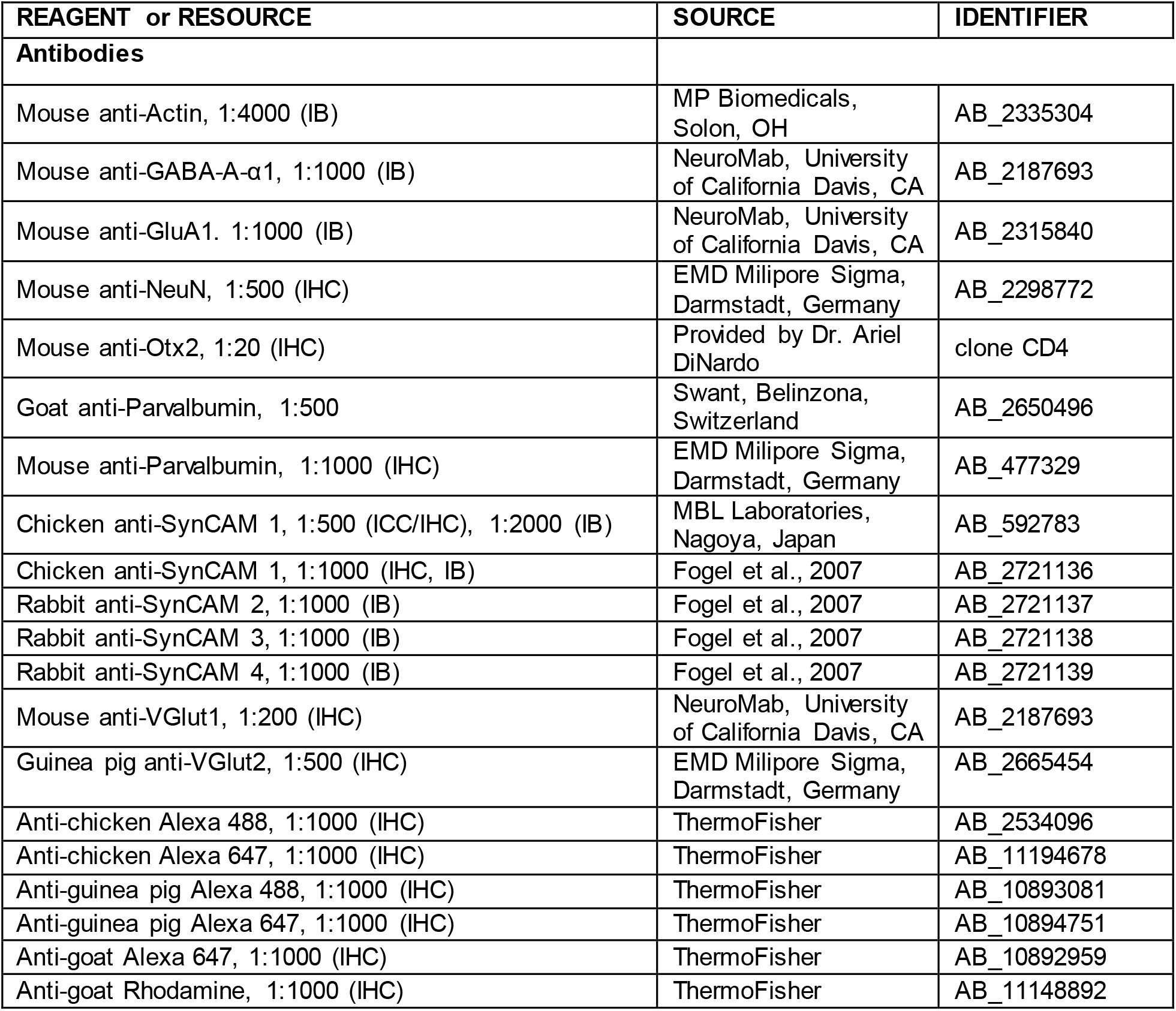
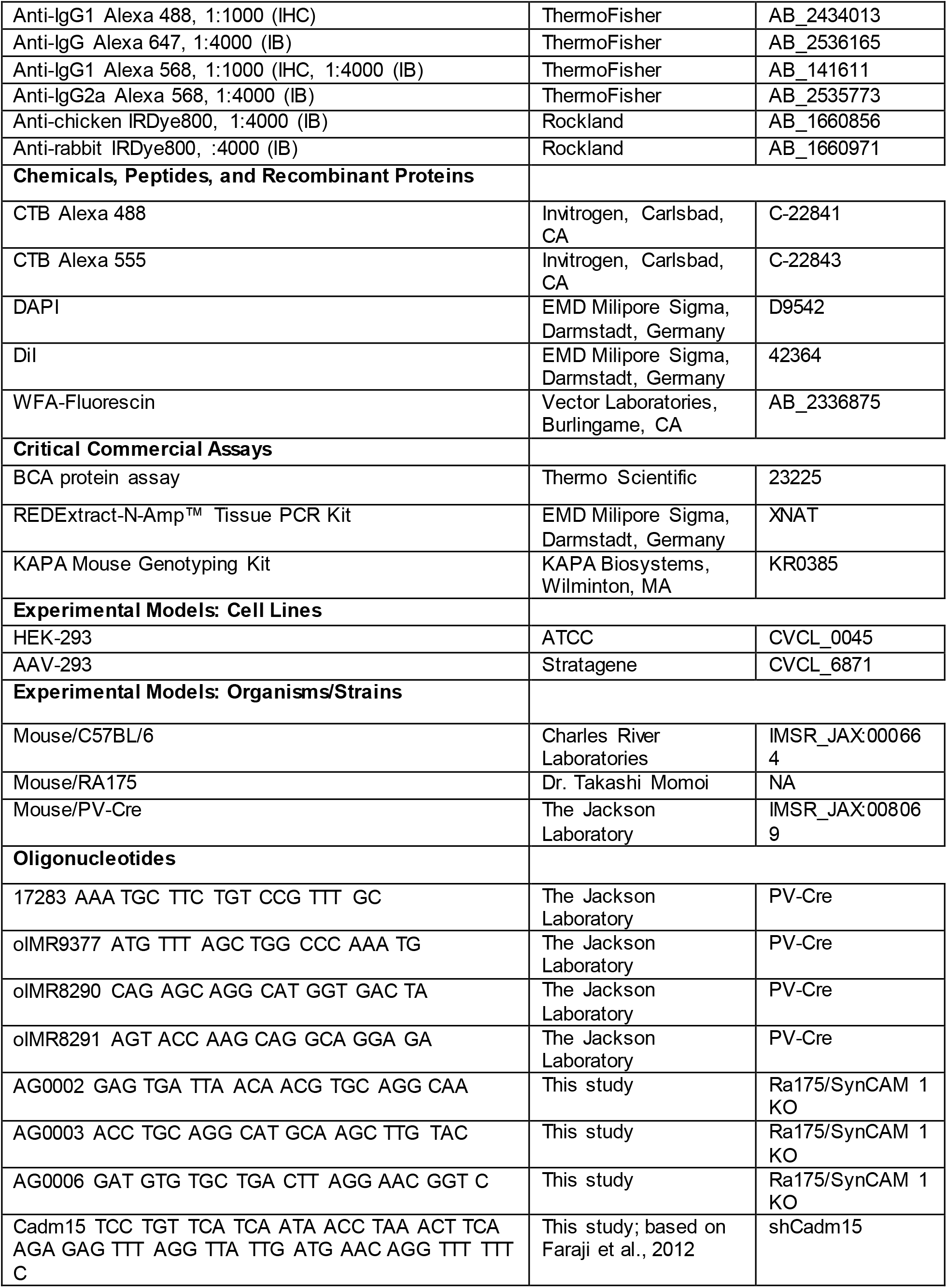
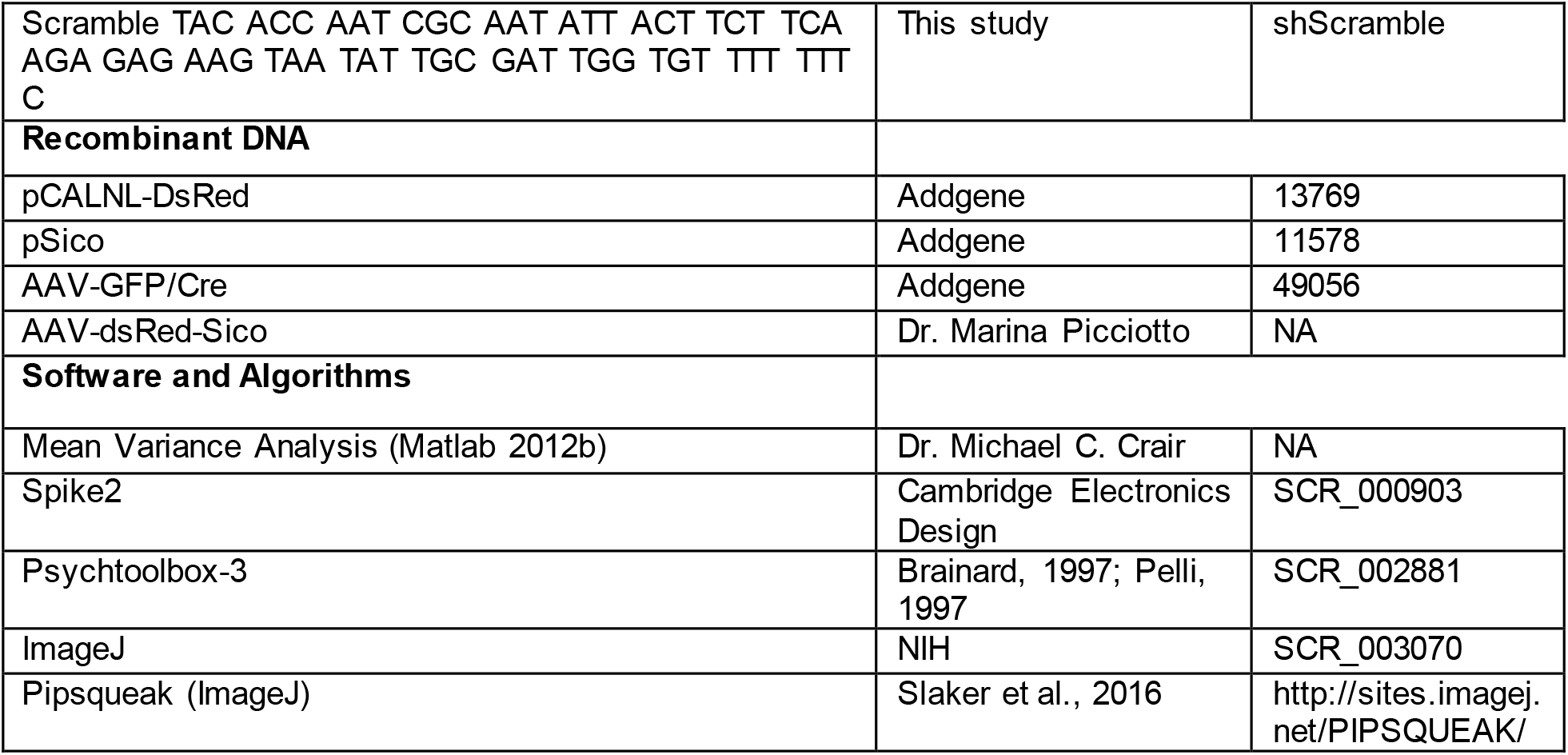
Key Resources Table.

